# Transcriptomic analysis of human induced pluripotent stem cell cardiomyocytes and myocardial tissue reveals novel role for miR-9-5p in hypertrophic cardiomyopathy

**DOI:** 10.64898/2026.09.07.749879

**Authors:** Doris M. T. Adao, Heta Lad, Mingyi Tang, Sophia G. Lopez, Filio Billia, Craig A. Simmons

## Abstract

Hypertrophic cardiomyopathy (HCM) is an inherited disease characterized by the thickening of the heart muscle. Mutations in genes encoding contractile proteins, such *MYBPC3* and *MYH7* account for over half of patients with familial HCM. MicroRNAs (miRNAs) are pivotal regulators of gene expression that may influence disease progression. Despite their potential significance, there is a scarcity of published data on miRNAs in HCM. Induced pluripotent stem cells (iPSCs) have emerged as valuable tools for modeling various cardiovascular diseases, including HCM. However, the miRNA expression profiles associated with HCM remain largely unexplored within iPSC models. Investigating the differential expression of miRNAs in HCM patients offers a promising avenue to identify potential therapeutic targets for individuals affected by this condition. In this study, we performed an unbiased screen using a microarray (NanoString Technologies) that contained 798 miRNAs to identify differentially expressed miRNAs in iPSC-cardiomyocytes (CM) and myocardial tissues derived from healthy donors and HCM patients with *MYBPC3* and *MYH7* mutations, identifying miR-9-5p as significantly downregulated in all HCM iPSC-CM and myectomy tissue compared to control. KEGG pathway analysis of gene targets using miRTarBase and g:Profiler revealed that miR-9-5p regulates genes in the epidermal growth factor receptor (EGFR) tyrosine kinase inhibitor resistance pathway, which is involved in hypercontractility associated with HCM. Modulation of miR-9-5p expression *in vitro* using mimics significantly reduced cardiomyocyte size and contractility of *MYBPC3* mutant iPSC-CMs whereas healthy control iPSC-CMs treated with antagomiRs exhibited hypertrophy. Our findings identify miR-9-5p as a regulator of cardiomyocyte hypertrophy and contractility, and suggest the miR-9-5p/EGFR tyrosine kinase inhibitor resistance axis as a potential therapeutic target to mitigate hypertrophy and hypercontractility exhibited in HCM.

## Introduction

Hypertrophic cardiomyopathy (HCM) is among the most common genetic disorders, with an estimated prevalence of about 1:200-500 in adults (Maron, 2002; Maron Barry J. et al., 1995; Semsarian et al., 2015). It is characterized by cardiomyocyte (CM) hypertrophy and myofibril disorganization at the cellular level (Marian & Braunwald, 2017) with over 100 mutations in more than 10 genes primarily encoding contractile sarcomere proteins. β-myosin heavy chain (MYH7) and cardiac myosin binding protein C (MYPBC3), are among the most commonly mutated genes, accounting for about half of familial HCM cases (Erdmann et al., 2003; Marian & Roberts, 2001).

Clinically, HCM is identified by left ventricular wall thickness (≥ 15 mm) without other secondary causes such as hypertension or changes in loading conditions (Erdmann et al., 2003). HCM manifests with a spectrum of phenotypes, ranging from mild conditions to aggressive forms of the disease with a high risk of sudden cardiac failure (Maron, 2018). HCM pathology shows marked heterogeneity, even within patients with mutations in the same gene (Page et al., 2012). Incomplete penetrance of the disease further complicates clinical decision-making. Despite this understanding, the mechanisms by which pathological hypertrophy occurs in response to mutations remain unclear. Recently, microRNAs (miRNAs) have gained attention as key regulators in disease progression due to their capacity to modulate gene expression by inhibiting messenger RNA (mRNA) translation or promoting mRNA degradation (Wahid et al., 2010). Their contribution to adverse cardiac remodeling in HCM has yet to be elucidated.

A key challenge in studying disease progression in genetically derived cardiac conditions is the scarcity of human models of disease, including biospecimens from the early stages of disease development. Typically, myectomy biospecimens from HCM patients are obtained in later stages, when patients develop severe myocardial hypertrophy causing obstruction of the left ventricular outflow tract. This leaves investigators with few opportunities to study disease progression. However, the advent of human induced pluripotent stem cells (iPSCs) provides an opportunity to overcome these limitations. iPSCs have the capacity to differentiate into various somatic cell types, including cardiomyocytes, carrying specific mutations associated with HCM (Li et al., 2022). This opens avenues for investigating how changes in miRNA expression contribute to the progression of HCM *in vitro*, and the possibility of using miRNA therapeutics to mitigate disease phenotypes.

With iPSC-cardiomyocytes (iPSC-CM) from patients with HCM harboring mutations in *MYBPC3* and *MYH7,* we investigated the differential expression of miRNAs and validated our findings using myectomy biospecimens. We identified miR-9-5p as significantly downregulated in HCM and the miR-9-5p regulated genes included the epidermal growth factor receptor (EGFR) tyrosine kinase inhibitor resistance pathway, a mechanism previously linked to impaired contractility and increased cardiomyocyte apoptosis in HCM. CM size and contractility were altered with the use of miR-9-5p mimics and inhibitors, highlighting the miR-9-5p-EGFR tyrosine kinase axis as a critical player in HCM pathology and a promising therapeutic target to mitigate disease.

## Results

### Patient cohort characteristics

Blood samples differentiated into iPSC-CM were donated by one male HCM patient with the *MYBPC3* mutation and one female HCM patient with the *MYH7* mutation (Supplementary Figure 1). One respective family member that was both genotype and phenotype negative also donated blood to serve as healthy controls. Despite having similar body mass index (BMI) (*MYBPC3* = 31.6 and *MYH7* = 34.1 kg/m^2^), and being younger in age at the time of donation (*MYBPC3* = 63.3 y and *MYH7* = 50.6 y), the *MYH7* iPSC-CM donor exhibited a larger left ventricular (LV) size (LV end-diastolic diameter 5.2 cm versus 3.7 cm; LV end-systolic diameter: 3.3 cm versus 2.5 cm), and LV mass index (110.0 g/m^2^ versus 67.4 g/m^2^), compared to the *MYBPC3* iPSC-CM donor. The LVEF was not significant different between the *MYH7* iPSC-CM and *MYBPC3* iPSC-CM donor (59.0% versus 62%; Table 1). It is difficult to define disease severity based this limited sample size.

**Table 1.** iPSC Donor Patient Characteristics (*N* = 2)

|  | *MYBPC3* (n = 1) | *MYH7* (n = 1) |
| --- | --- | --- |
| **Mutation** | p.Gln1233Ter | p.Ala355Ser |
| **Characteristics** |  |  |
| Age at Donation | 63.3 | 50.6 |
| Male | 0 | 1 |
| Female | 1 | 0 |
| Height (cm) | 164 | 182 |
| Weight (kg) | 85 | 113 |
| BMI (kg/m<sup>2</sup>) | 31.6 | 34.1 |
| **Echocardiography Data** |  |  |
| LV End-Diastolic Diameter (cm) | 3.7 | 5.2 |
| LV End-Systolic Diameter (cm) | 2.5 | 3.3 |
| LV Ejection Fraction (%) | 62.0 | 59.0 |
| LV Mass (g) | 129.3 | 254.0 |
| LV Mass Index (g/m<sup>2</sup>) | 67.4 | 110.0 |

Data represents echocardiography measurements at time of blood donation later used for iPSC-CM differentiation. BMI = Body mass index; LV = Left ventricular.

Myocardial tissue samples were collected from HCM patients with *MYBPC3* (n = 4) and *MYH7* (n = 3) mutations who underwent surgical myectomy. Healthy control myocardial tissues were obtained from individuals whose hearts were not utilized for heart transplantation (n = 3). There were no significant differences in age and body mass index between donor cohorts at the time of myectomy procedures (Table 2). Myectomy tissue patients with *MYH7* mutations had significantly increased interventricular septal thickness at end diastole compared to the control group (2.1 ± 0.1 cm versus 1.0 ± 0.1 cm; p = 0.03), but no other statistically significant differences were present on echocardiographic assessment (Table 2). This suggests that our *MYBPC3* and *MYH7* donor groups are grossly phenotypically similar in cardiac structure and function.

**Table 2.** Myectomy Tissue Donor Patient Characteristics (*N* = 10)

|  | *MYBPC3* (n = 4) | *MYH7* (n = 3) | Control (n = 3) | *P* Value |
| --- | --- | --- | --- | --- |
| **Characteristics** |  |  |  |  |
| Age at Myectomy | 54.1 ± 17.1 | 54.7 ± 25.0 | 58.7 ± 4.7 | 0.67 |
| Male | 2 | 1 | 2 | . |
| Female | 2 | 2 | 1 | . |
| Height (cm) | 166.0 ± 7.8 | 165.0 ± 8.9 | . | 0.87 |
| Weight (kg) | 85.8 ± 35.0 | 92.0 ± 3.6 | . | 0.51 |
| BMI (kg/m<sup>2</sup>) | 30.5 ± 9.6 | 34.1 ± 4.8 | . | 0.35 |
| **Echocardiography Data** |  |  |  |  |
| LV End-Diastolic Diameter (cm) | 4.3 ± 0.2 | 4.3 ± 0.5 | 4.6 ± 0.8 | 0.61 |
| LV End-Systolic Diameter (cm) | 2.6 ± 0.6 | 2.4 ± 0.2 | 2.9 ± 0.5 | 0.45 |
| LV Ejection Fraction (%) | 73.0 ± 10.0 | 49.7 ± 29.5 | 64.5 ± 8.5 | 0.27 |
| LV Mass (g) | 244.3 ± 81.0 | 308.6 ± 20.3 | 114 | 0.11 |
| LV Mass Index (g/m<sup>2</sup>) | 131.3 ± 39.2 | 154.2 ± 5.2 | 84.25 ± 25.8 | 0.11 |
| Interventricular Septal Thickness End Diastole (cm) | 1.8 ± 0.7 | 2.1 ± 0.1* | 1.0 ± 0.1 | 0.03 |
| LV Posterior Wall Diameter (cm) | 1.1 ± 0.1 | 1.0 ± 0.2 | 1.0 ± 0.02 | 0.71 |
| LV Maximal Wall Thickness (mm) | 22.5 ± 4.5 | 22.7 ± 1.5 | . | 0.95 |

Data represents echocardiography measurements before cardiac myectomy procedure.
Statistical analysis was conducted using one-way ANOVA with Tukey's multiple comparisons
test. (\* $p \leq 0.05$ ). Data is presented as mean ± SD. BMI = Body mass index; LV = Left ventricular.

### Differential miRNA expression of in iPSC-CM versus myectomy tissue

Total RNA was isolated from *MYBPC3* and *MYH7* iPSC-CM and myectomy tissue samples and sent for NanoString microarray analysis. Raw miRNA counts were normalized using nSolver (v.04) and fold-change expression was calculated between mutant and control samples. In *MYBPC3* iPSC-CMs, 26 miRNAs were significantly downregulated and 23 were significantly upregulated in mutants compared to control (Figure 1A). In *MYBPC3* mutant myectomy tissues, 55 miRNAs are significantly downregulated and 23 were significantly upregulated (Figure 1B). In *MYH7* iPSC-CMs, 38 miRNAs were significantly downregulated and 29 were significantly upregulated (Figure 1C), and 24 miRNAs were downregulated and 12 were significantly upregulated in *MYH7* myectomy tissues (Figure 1D). When comparing the expression of significantly differentially expressed miRNAs, the overlap was greater within the same sample type (myectomy versus iPS-CM) than between different sample types that shared the same gene mutation (e.g., iPSC-CM and tissue samples both with *MYBPC3* or *MYH7* mutations). In addition, there were 10 miRNAs that were concordantly differentially expressed in *MYBPC3* samples 6 miRNAs for *MYH7* samples (Figure 1E). Interestingly, one miRNA, miR-9-5p, was significantly downregulated among all the samples (Figure 1E).

**Figure 1.**
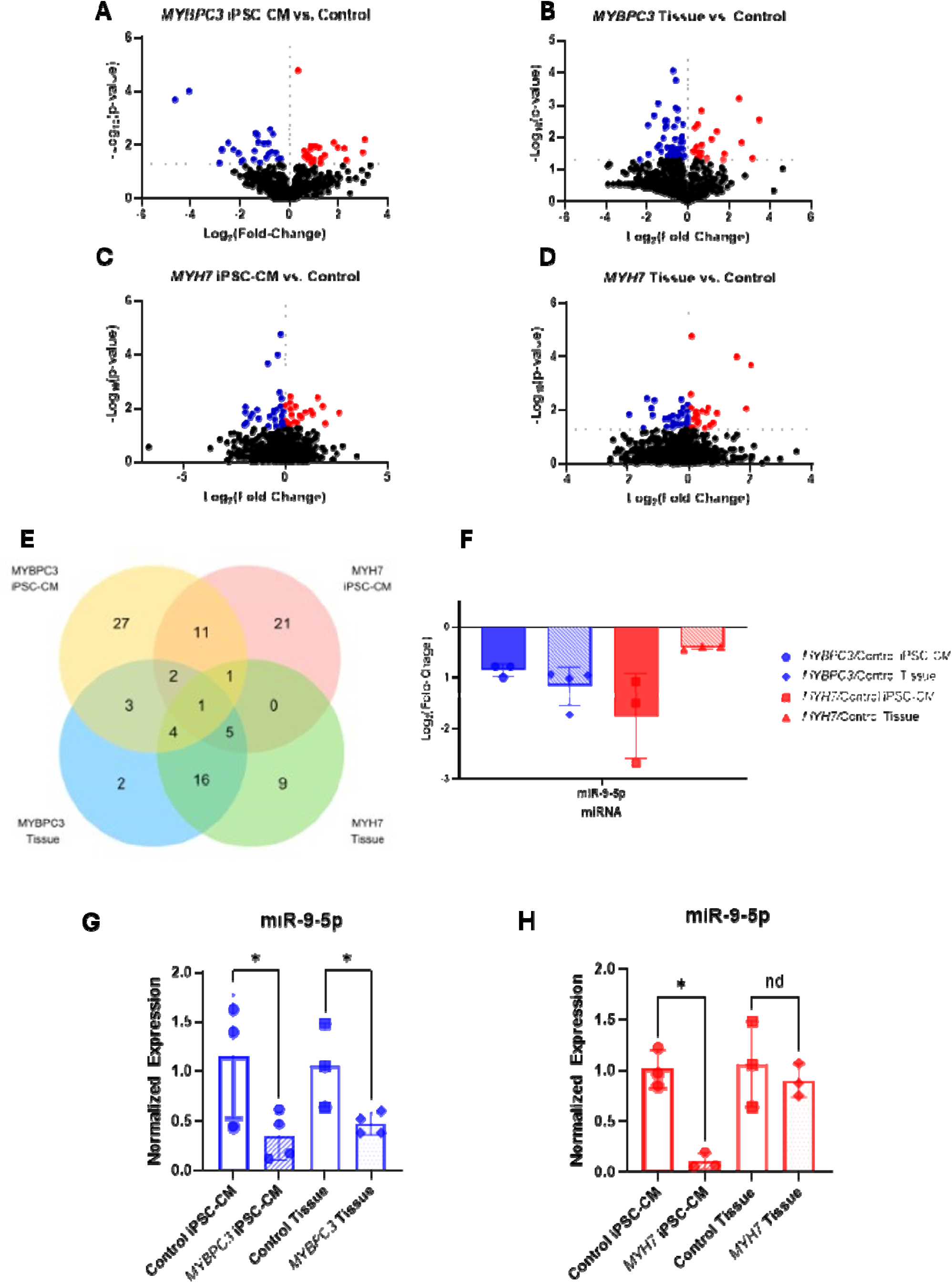
Differentially expressed miRNAs in iPSC-CM and myectomy tissue of HCM patients with *MYBPC3* and *MYH7* mutations. NanoString microarray analysis revealed significantly upregulated and downregulated miRNAs in **(A)** *MYBPC3* iPSC-CM (n = 3) vs. *MYBPC3* control iPSC-CM (n = 3), **(B)** *MYBPC3* tissue (n = 4) vs. control tissue (n = 3), **(C)** *MYH7* iPSC-CM (n = 3) vs. *MYH7* control iPSC-CM (n = 3), and **(D)** *MYH7* tissue (n = 3) vs. control tissue (n = 3). **(E)** Venn diagram of miRNA overlaps between iPSC-CM and myectomy samples. **(F)** Log_2_(Fold-Change) expression was determined for miR-9-5p in iPSC-CM and tissue samples using NanoString microarray analysis. RT-qPCR analysis of miRNAs in both iPSC-CM and myectomy tissues of **(G)** *MYBPC3* mutant and control groups and **(H)** *MYH7* mutant and control groups. Data is presented as mean ± SD. (* p ≤ 0.05).

When RT-qPCR was performed to validate these findings, miR-9-5p were significantly downregulated in *MYBPC3* iPSC-CM and myectomy tissue samples (Figure 1G), and significantly downregulated in *MYH7* iPSC-CM (Figure 1H).

### EGFR tyrosine kinase inhibitor resistance pathways is targeted by miR-9-5p

Given the consistent downregulation of miR-9-5p in mutant samples, miRTarBase (v9.0) was used to curate a list of miR-9-5p targets supported by strong experimental evidence (Supplementary Table 1). This gene set was subsequently analyzed using g:Profiler to assess pathway enrichment (Figure 2A), and protein–protein interactions were visualized using STRING (Figure 2B). Gene Ontology (GO) analysis revealed significant enrichment of terms such as enzyme binding, transmembrane receptor protein kinase activity, and regulation of cell development, with chromatin and nucleoplasm identified as prominent cellular components (Figure 2C). KEGG pathway analysis further identified microRNAs in cancer and EGFR tyrosine kinase inhibitor resistance as the most significantly enriched pathways (Figure 2D). The EGFR pathway has been implicated in the regulation of cardiomyocyte contractility and disease progression in HCM (Algül et al., 2023; Ewoldt et al., 2024), making it a pathway of interest for further investigation.

**Figure 2.**
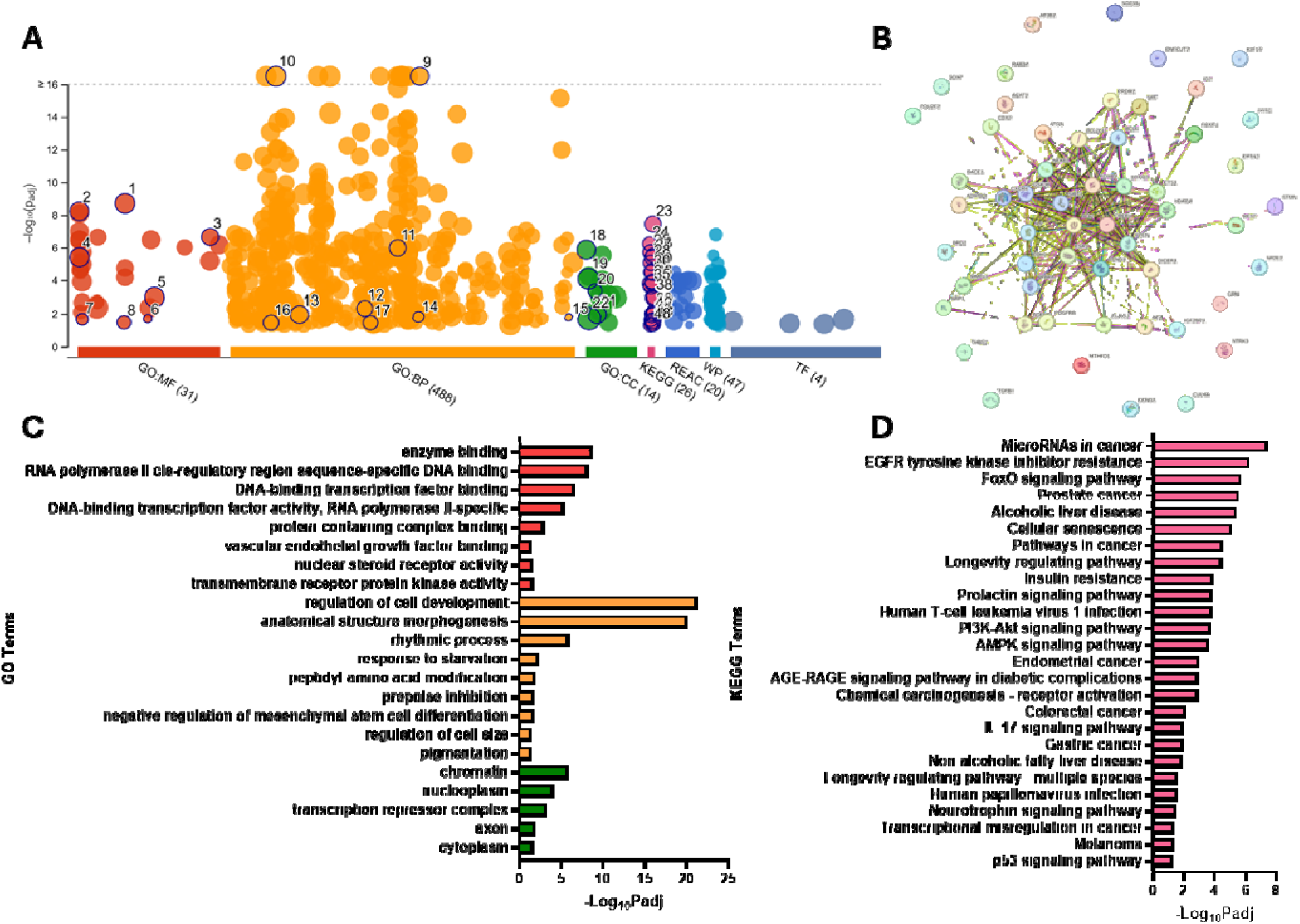
Pathway analysis of genes regulated by miR-9-5p using g:Profiler. Gene targets that were experimentally validated to interact with miR-9-5p were generated using miRTarBase (v.9.0). **(A)** A Manhattan plot of Top GO terms and KEGG terms that are significantly enriched by miR-9-5p gene targets are highlighted. **(B)** An interactome was generated using STRING (v.12.0) to visually map interactions between gene targets. **(C)** Top 22 GO Terms segmented by molecular function (red), biological processes (orange), and cellular component (green). **(D)** Top 27 KEGG pathways, including microRNAs in cancer and EGFR tyrosine kinase inhibitor resistance.

Further investigation of g:Profiler analysis revealed several miR-9-5p-regulated genes in the EGFR tyrosine kinase inhibitor (TKI) resistance pathway, including *FOXO3*, *NF1*, *PTEN*, *BCL2L11* (*Bim*), *BCL2*, *IL6*, *GSK3B*, and *PDGFRB* (Supplementary Table 2). We hypothesized that these gene targets would be upregulated in HCM since miR-9-5p is significantly downregulated in disease states. Indeed, RT-qPCR analysis identified several target genes that were significantly upregulated in a mutation– and tissue-specific manner, implicating pathways relevant to hypertrophic cardiomyopathy (Figure 3). In *MYBPC3* iPSC-CMs, upregulation of *FOXO3, NF1, PTEN*, and *GSK3B* suggests activation of stress-responsive and anti-hypertrophic signaling pathways that may represent compensatory mechanisms in response to early hypercontractile stress (Figure 3A). In *MYBPC3* myectomy tissue, increased *NF1* and *BCL2* expression indicates sustained modulation of growth factor signaling and enhanced pro-survival signaling in diseased myocardium (Figure 3C). In *MYH7* models, upregulation of *BCL2L11* in iPSC-CMs points to increased apoptotic susceptibility during early disease stages (Figure 3B), while elevated *BCL2* expression in *MYH7* myectomy tissue suggests a shift toward apoptosis resistance in advanced disease (Figure 3D). Collectively, the upregulation of these genes highlights dysregulation of survival, stress adaptation, and growth signaling pathways that are central to HCM pathogenesis and progression (Figure 3E).

**Figure 3.**
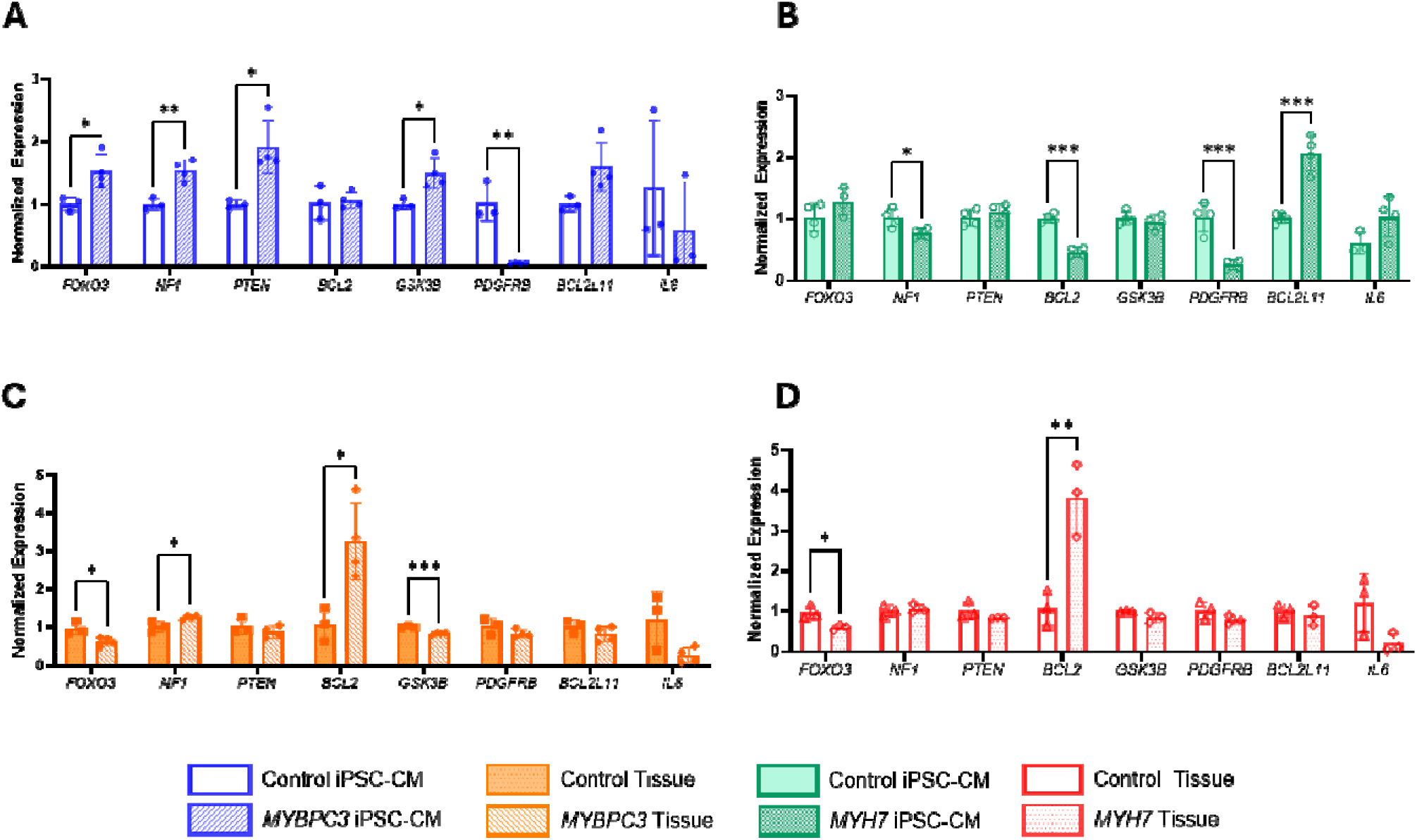
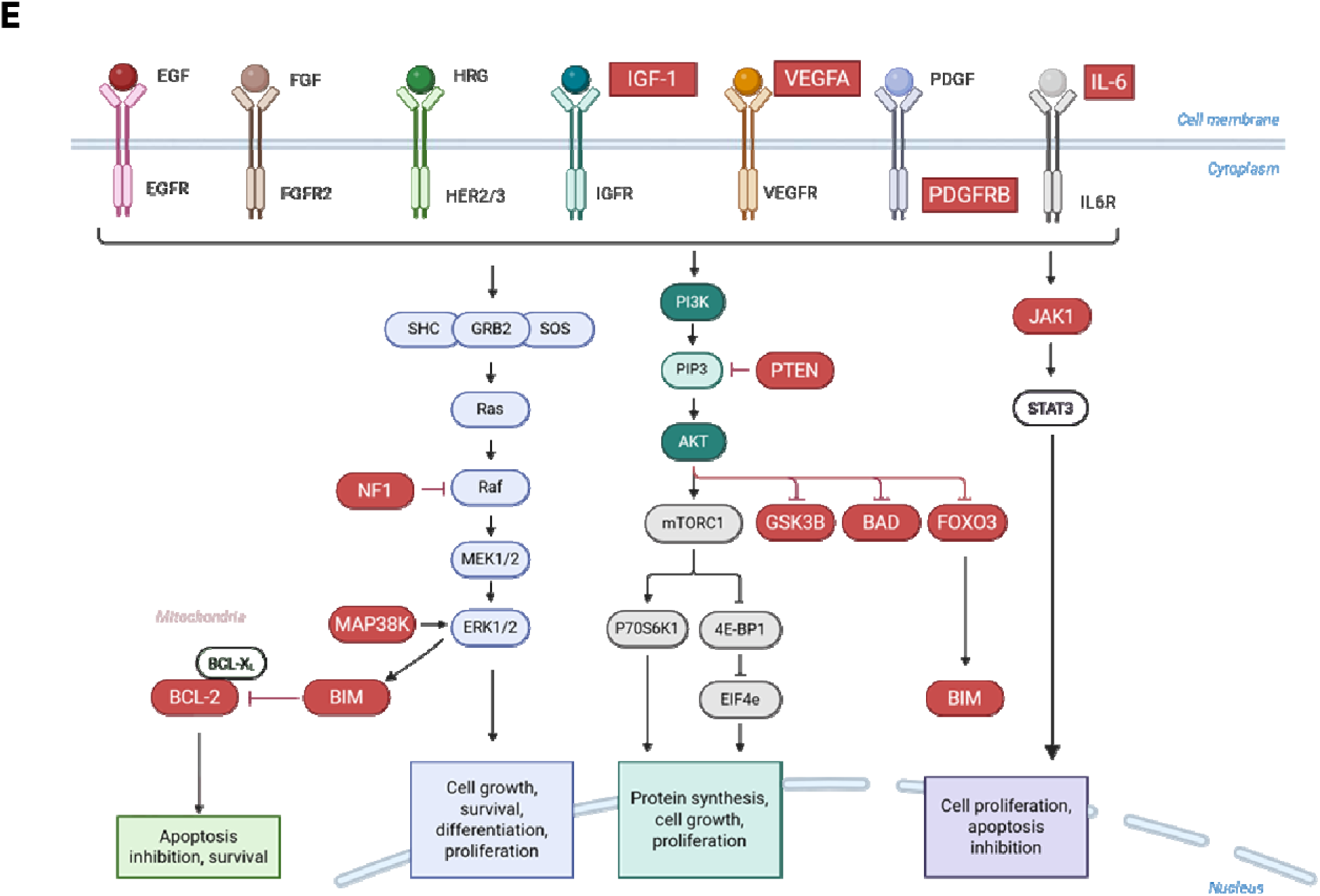
RT-qPCR validation of genes in EGFR tyrosine kinase inhibitor resistance pathway regulated by miR-9-5p. Gene expression was normalized using the –2ΔΔCt method and normalized to the 5S rRNA housekeeping gene set. **(A)** Represents gene terms significantly differentially expressed in *MYBPC3* iPSC-CM, **(B)** *MYBPC3* tissue, **(C)** *MYH7* iPSC-CM, and **(D)** *MYH7* tissue. Differences between mutant and control groups were determined using Student’s t-test (Graphpad Prism, v. 10.4.1). Data is presented as mean ± SD (* p ≤ 0.05, ** p ≤ 0.01, *** p ≤ 0.001). **(E)** Visual representation of gene terms regulated by miR-9-5p in the EGFR tyrosine kinase inhibitor resistance pathway highlighted in red.

**Figure 4.**
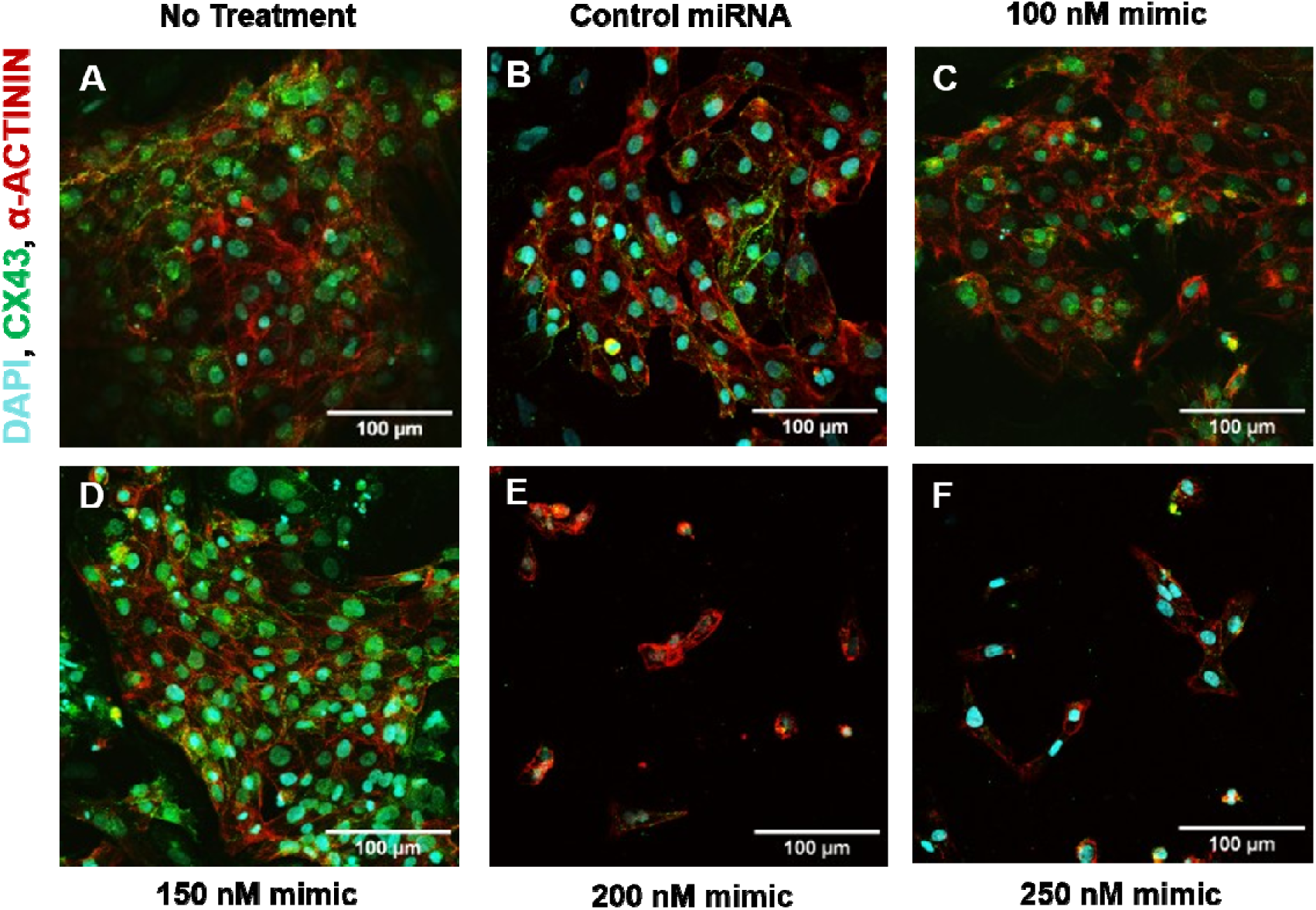
Immunofluorescence images of *MYBPC3* mutant iPSC-CM. Cells were stained with DAPI, Connexin-43 (CX43), and α-actinin. *MYBPC3* mutant iPSC-CM were treated with different dose levels of miR-9-5p mimics **(A)** No Treatment, **(B)** Control miRNA, **(C)** 100 nM, **(D)** 150 nM, **(E)** 200 nM, and **(F)** 250 nM mimics.

### miR-9-5p mimics treatment alters the hypertrophic phenotype in vitro

To investigate the role of miR-9-5p *in vitro*, transfection experiments were conducted on *MYBPC3*-mutant iPSC-CMs using miR-9-5p mimics to upregulate miR-9-5p expression with the goal of mitigating disease-associated phenotypes. *MYBPC3*-mutant iPSC-CMs were treated with miR-9-5p mimics at varying dose levels, and there were no differences in the surface area measured (p = 0.0743). However, there was a significant decrease in surface area when No Treatment and Control miRNA groups were compared to the various concentrations of the miR-9-5p mimics (p < 0.0001) (Figure 5A). Collectively, these results demonstrate that miR-9-5p mimic transfection effectively reduced CM surface area in *MYBPC3*-mutant iPSC-CMs, indicating a reversal of the hypertrophic phenotype in this disease model.

**Figure 5.**
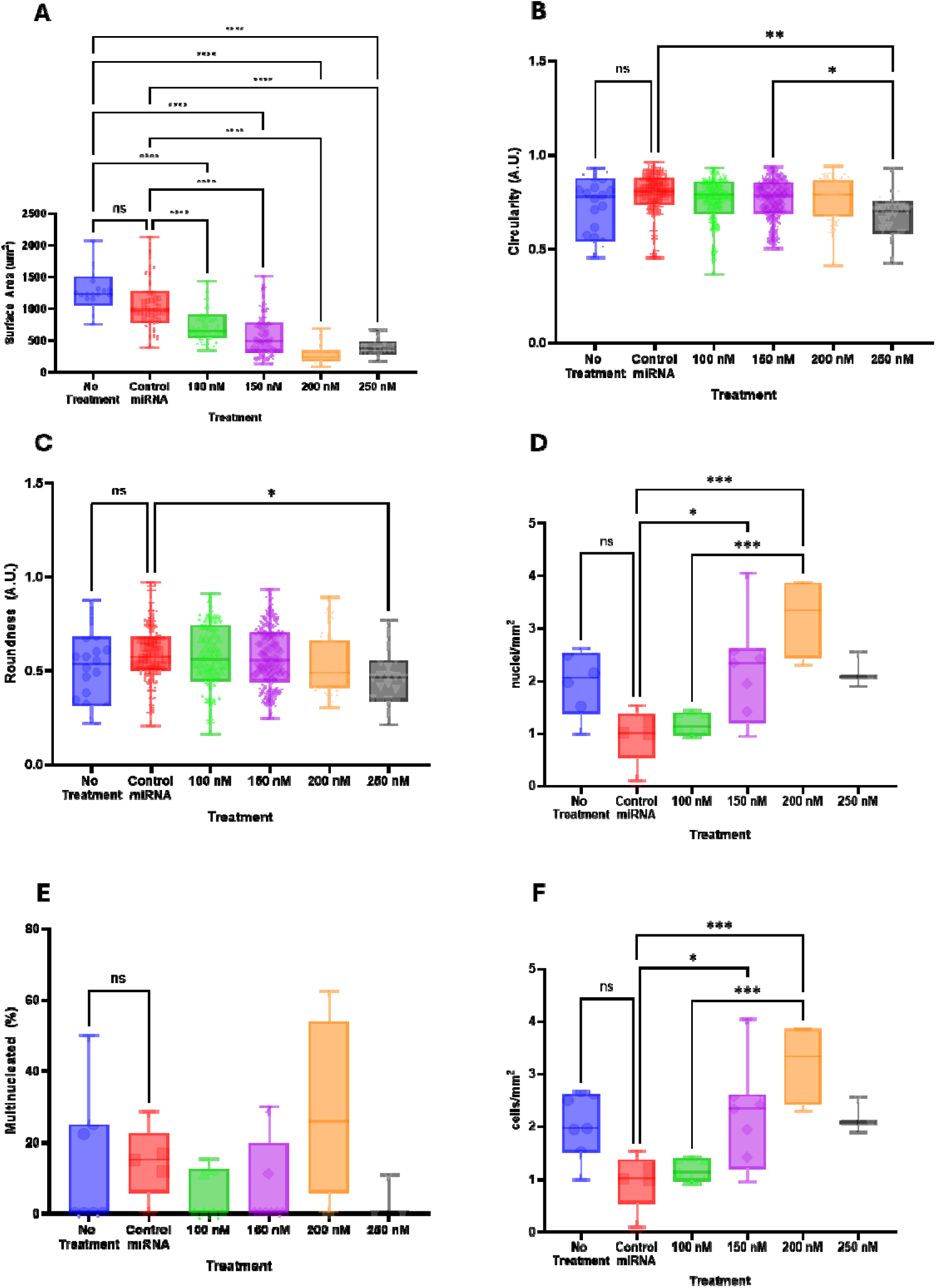
Quantification of *MYBPC3* mutant iPSC-CM morphology after treatment with different dose levels of miR-9-5p mimics. To determine cardiomyocyte **(A)** Surface area, **(B)** Circularity, and **(C)** Roundness, iPSC-CM with clear cell borders were identified across multiple wells, imaged, and traced using ImageJ in each of the following conditions: No treatment (n = 24), Control miRNA (n = 52), 100 nM (n = 65), 150 nM (n = 115), 200 nM (n = 34) and 250 nM (n = 23). **(D)** Nuclear density, **(E)** Multinucleation, and **(F)** Cell density, were derived from a series of immunofluorescence images. Statistical comparisons between groups were calculated using one-way ANOVA with Tukey’s multiple comparisons test (* p ≤ 0.05, ** p ≤ 0.01, *** p ≤ 0.001, **** p ≤ 0.001). Data is presented as mean ± min and max data points.

To further characterize whether miR-9-5p upregulation alters CM morphology beyond changes in cell size, we quantified circularity and roundness of *MYBPC3*-mutant iPSC-CMs across treatment groups, with a score of 1.0 indicating a perfect circle and values approaching 0.0 indicating increasingly elongated and polarized shapes. Circularity reflects shape complexity by assessing changes in perimeter while area remains constant, whereas roundness evaluates elongation by comparing the ratio between CM length and width. A significant difference in iPSC-CM circularity was observed between the Control miRNA group (0.79 ± 0.12 A.U.) and the 250 nM-treated group (0.68 ± 0.12 A.U.; p = 0.0035), as well as between the 150 nM group (0.76 ± 0.11 A.U.) and the 250 nM-treated group (0.68 ± 0.12 A.U.; p = 0.0424) (Figure 5B). In contrast, significant differences in iPSC-CM roundness were only observed between the Control miRNA group (0.59 ± 0.16 A.U.) and the 250 nM-treated group (0.46 ± 0.15 A.U.; p = 0.0292) (Figure 5C). Together, these findings suggest that high-dose miR-9-5p upregulation could induce modest changes in CM shape, however, this could likely be a toxic effect at high doses of mimic treatment.

To determine whether miR-9-5p upregulation influences CM density and cellular organization in addition to cell size, nuclear and cell density were quantified across treatment groups. Nuclear density was observed by counting number of nuclei over the total α-actinin-positive area. There was no significant difference in nuclei density between the No Treatment and Control miRNA conditions (p = 0.1895). Compared to the Control miRNA-treated group, there were significant increases in nuclei density in iPSC-CM treated with 150 nM of miR-9-5p mimics (2.15 ± 0.97 nuclei/mm^2^; p = 0.0435) and 200 nM of miR-9-5p mimics (3.21 ± 0.78 nuclei/mm^2^; p = 0.0004) (Figure 5D). To calculate cell density, multinucleated iPSC-CM were identified from iPSC-CM with clear borders (Figure 5E), and total number of cells was calculated assuming an average of 2 nuclei per multinucleated cell. Once again, there were no differences in cell density between No Treatment and Control miRNA-treated groups, but compared to the Control miRNA-treated group, there was a significant increase in cell density in the 150 nM-treated group (2.15 ± 0.97 cells/mm^2^; p = 0.0422), and 200 nM-treated group (3.21 ± 0.77 cells/mm^2^; p = 0.0004) (Figure 5F). Together, these results indicate that miR-9-5p upregulation increases nuclear and cell density in *MYBPC3*-mutant iPSC-CMs, consistent with reduced CM hypertrophy and cellular organization rather than overt proliferative expansion. Therefore, mimic-mediated upregulation of miR-9-5p in *MYBPC3*-mutant iPSC-CMs reduced CM size, increased nuclear and cell density, and promoted shape changes consistent with attenuation of disease-associated hypertrophic remodeling.

### Treatment using miR-9-5p antagomiRs enhanced the hypertrophic response in vitro

In an effort to recapitulate disease-associated hypertrophic remodeling observed in HCM, miR-9-5p expression was downregulated in healthy control iPSC-CMs using miR-9-5p antagomiRs. Control iPSC-CMs were treated with miR-9-5p antagomiRs at various doses, and surface area was measured (Figure 6). There was no significant difference in cross-sectional area between the No Treatment and Control miRNA groups (p = 0.8376), but there was a significant increase in iPSC-CM surface area when treated with miR-9-5p antagomiRs at 25 nM (p < 0.0001), 50 nM (p < 0.0001), 75 nM (p < 0.0002), and 100 nM (p < 0.0001) compared to the No Treatment group (Figure 7A). When comparing surface area between the Control miRNA group and miR-9-5p antagomiR-treated groups, significant increases were observed at 25 nM (p = 0.0004) and 50 nM (p < 0.0001) (Figure 7A). Collectively, these findings demonstrate that miR-9-5p downregulation induces CM hypertrophy in healthy control iPSC-CMs, recapitulating a key cellular feature of HCM and supporting a role for miR-9-5p in restraining pathological CM growth.

**Figure 6.**
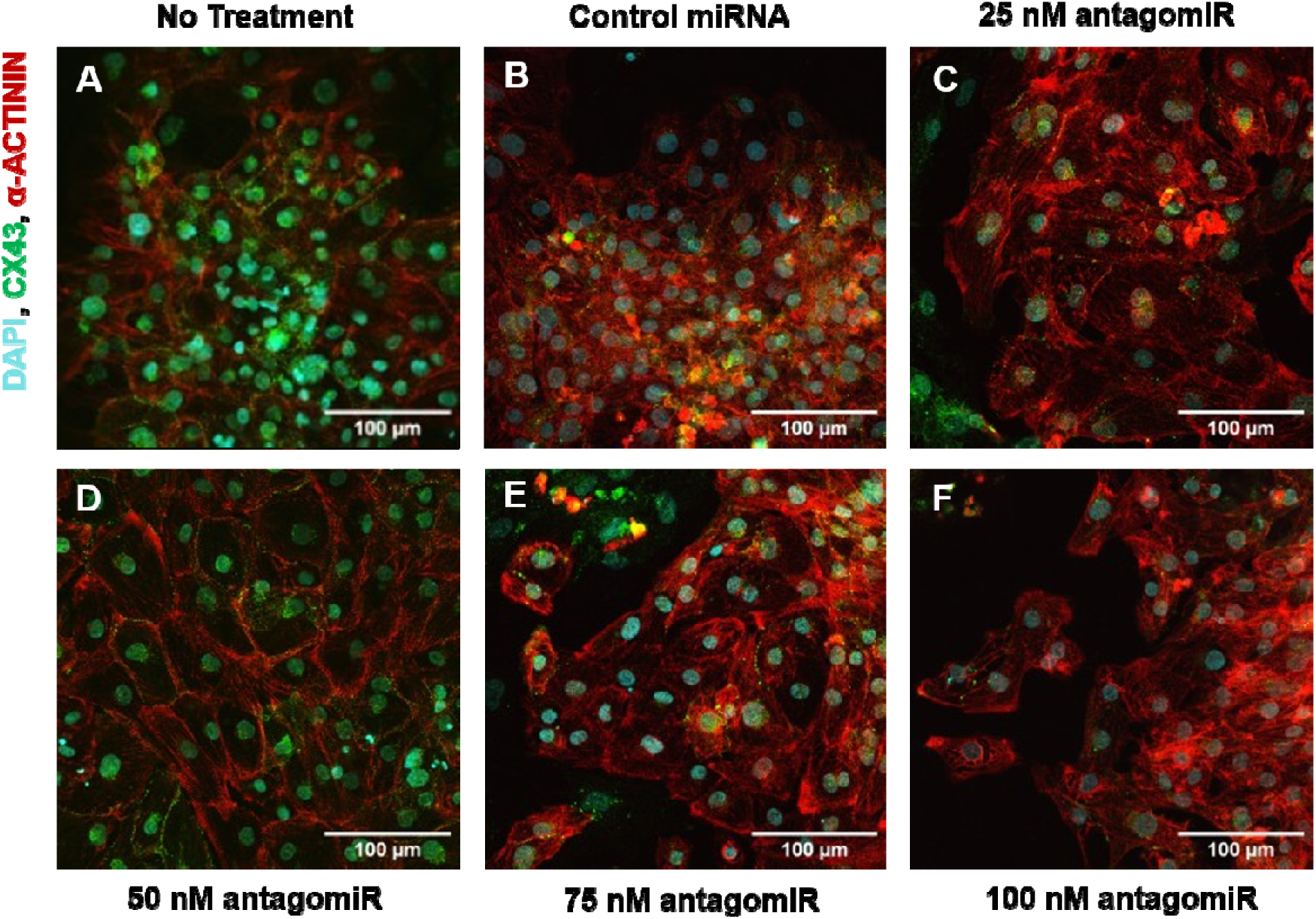
Immunofluorescence images of healthy control iPSC-CM. Cells were stained with DAPI, Connexin-43 (CX43), and α-actinin. Control iPSC-CM were treated with different dose levels of miR-9-5p antagomiRs: **(A)** No Treatment, **(B)** Control miRNA, **(C)** 25 nM, **(D)** 50 nM, **(E)** 75 nM, and **(F)** 100 nM antagomiR.

**Figure 7.**
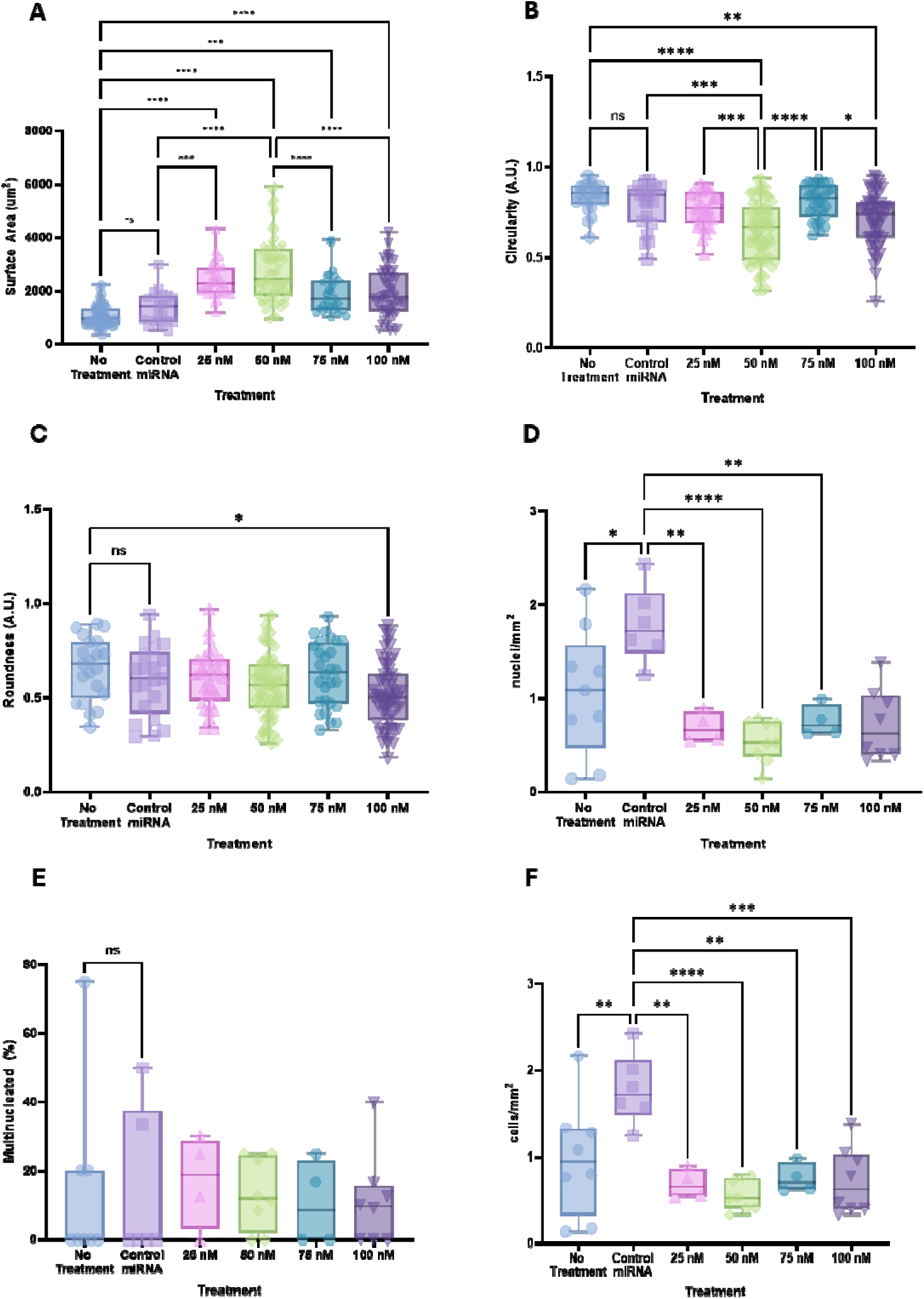
Quantification of healthy control iPSC-CM morphology after treatment with different dose levels of miR-9-5p antagomiRs. To determine cardiomyocyte **(A)** Surface area, **(B)** Circularity, and **(C)** Roundness, iPSC-CM with clear cell borders were identified across multiple wells, imaged, and traced using ImageJ in each of the following conditions: **(A)** No treatment (n = 65), **(B)** Control miRNA (n = 21), **(C)** 25 nM (n = 32), **(D)** 50 nM (n = 67), **(E)** 75 nM (n = 28) and **(F)** 100 nM (n = 63). **(D)** Nuclear density, **(E)** Multinucleation, and **(F)** Cell density, were derived from a series of immunofluorescence images. Statistical comparisons between groups were calculated using one-way ANOVA with Tukey’s multiple comparisons test (* p ≤ 0.05, ** p ≤ 0.01, *** p ≤ 0.001, **** p ≤ 0.001). Data is presented as mean ± min and max data points.

To assess whether miR-9-5p downregulation induces CM shape changes associated with pathological remodeling observed in HCM, circularity and roundness were quantified across antagomiR treatment groups. Circularity was not significantly different between the No Treatment and Control miRNA-treated groups. However, there were significant decreases in circularity between the No Treatment and 50 nM group (0.65 ± 0.17 A.U.; p < 0.0001), as well as the 100 nM group (0.71 ± 0.14 A.U.; p = 0.0061). The Control miRNA-treated group was also less circular than the 50 nM antagomiR-treated group (0.65 ± 0.17 A.U.; p = 0.0007). The iPSC-CM treated with 50 nM of miR-9-5p antagomiR (0.65 ± 0.17 A.U.) was also significantly less circular than 25 nM-treated (0.76 ± 0.10 A.U; p = 0.0007) and 75 nM-treated groups (0.81 ± 0.09 A.U; p = < 0.0001). The 100 nM-treated group (0.71 ± 0.14 A.U) had lower circularity compared to the 75 nM-treated group (p = 0.0289) (Figure 7B). When comparing roundness, there was a significant difference between the No Treatment Group (0.65 ± 0.17 A.U.) and the 100 nM miR-9-5p antagomiR-treated group (0.52 ± 0.17 A.U.; p = 0.0176) (Figure 7C). Together, these findings indicate that miR-9-5p downregulation induces dose-dependent alterations in cardiomyocyte geometry that are consistent with pathological hypertrophic remodeling associated with HCM, characterized by increased cell size and altered cell shape.

### Treatment of miR-9-5p mimics and antagomirs alters the contractile phenotype of HCM in vitro

Hypercontractility and accelerated contraction kinetics are hallmark features of HCM, particularly in CM harboring sarcomeric mutations such as *MYBPC3*. To determine whether modulation of miR-9-5p influences these disease-associated contractile abnormalities, iPSC-CM contractility was assessed following transfection with miR-9-5p mimics and antagomiRs using particle image velocimetry (PIV), In *MYBPC3* mutant iPSC-CMs (Figure 8A), treatment with miR-9-5p mimics significantly reduced the maximum displacement during peak contraction in cells treated at concentrations of 100 nM (1.1 ± 0.3 μm; p = 0.0129) and 150 nM (1.0 ± 0.2 μm; p = 0.0030), compared to No Treatment (1.5 ± 0.5 μm) (Figure 8B). Similarly, contraction cycle length was significantly shortened in iPSC-CMs treated with 100 nM (1100 ± 260 ms; p = 0.0003) and 150 nM (1100 ± 210 ms; p < 0.0001) of miR-9-5p mimics, compared to untreated *MYBPC3* iPSC-CMs (1700 ± 430 ms) (Figure 8C). Time to peak contraction was also significantly reduced with 100 nM (530 ± 140 ms; p = 0.0009) and 150 nM (500 ± 93 ms; p = 0.0004) treatments, relative to the No Treatment group (1100 ± 600 ms) (Figure 8D). However, no significant changes were observed in the relaxation time (Figure 8E). These results demonstrate that miR-9-5p upregulation modulates contractile magnitude and kinetics in *MYBPC3* mutant iPSC-CMs in a manner consistent with attenuation of HCM-associated hypercontractility, supporting a functional role for miR-9-5p in regulating disease-relevant contractile phenotypes.

**Figure 8.**
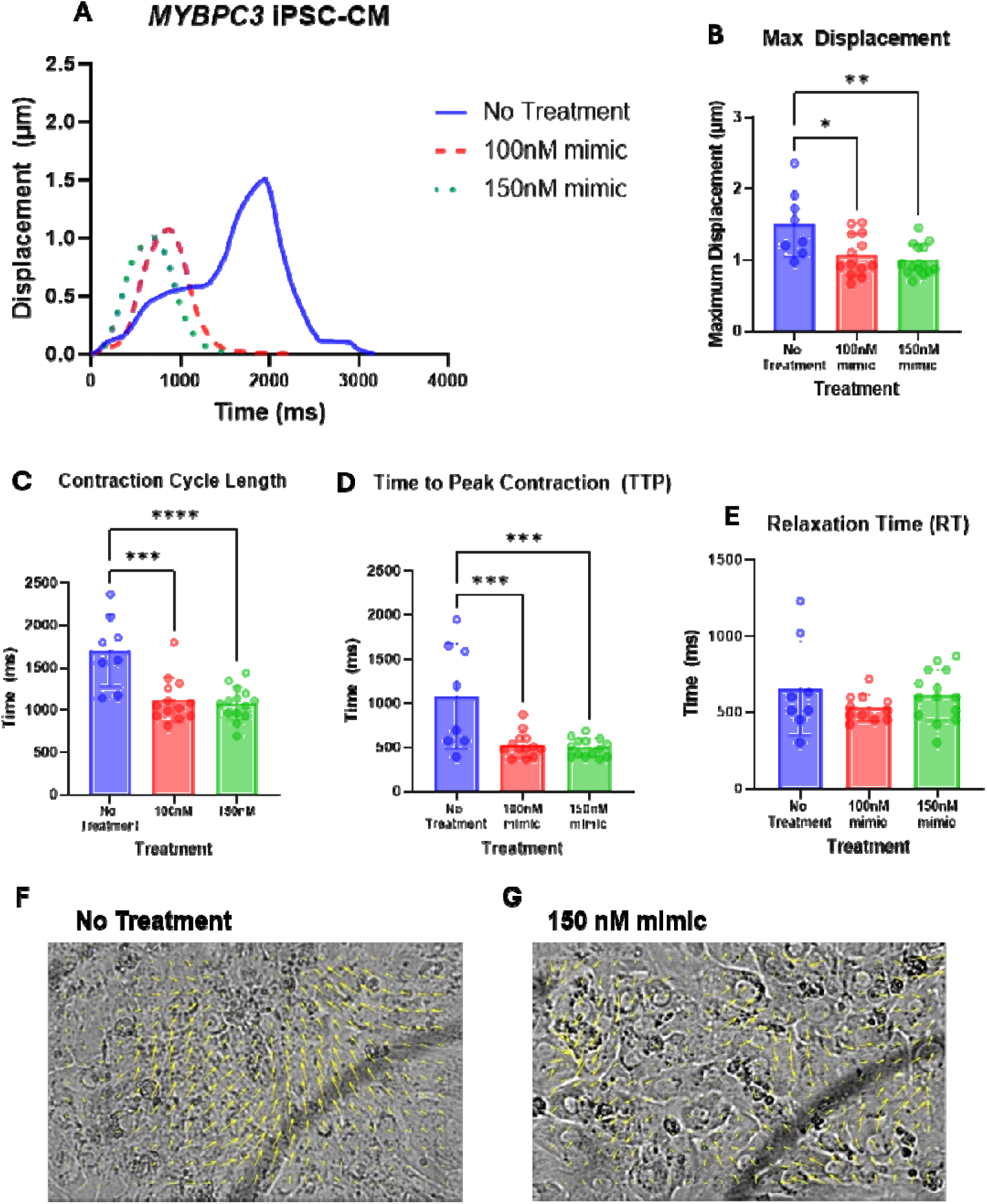
Contractility of *MYBPC3* iPSC-CM observed using Particle Image Velocimetry (PIV). Images were obtained using phase contrast microscopy, and particle displacement magnitude (µm) was measured over time (ms). **(A)** Representative contraction curves of *MYBPC3* mutant iPSC-CMs: No Treatment (n = 8) or treated with 100 nM (n = 13) or 150 nM (n = 14) of miR-9-5p mimics. Quantification of contractility parameters in *MYBPC3* mutant iPSC-CM: **(B)** Maximum displacement, **(C)** Contraction cycle length, **(D)** Time to peak contraction (TTP) **(E)** Relaxation time. **(F)** Vector plot arrows (in yellow) depicting contraction direction of untreated *MYBPC3* iPSC-CM and **(G)** *MYBPC3* iPSC-CM treated with 150 nM of miR-9-5p mimics. Videos can be accessed at https://doi.org/10.5683/SP3/GUBIRH. Statistical analysis was performed using one-way ANOVA with Tukey’s multiple comparisons test (* p ≤ 0.05, ** p ≤ 0.01, *** p ≤ 0.001, **** p ≤ 0.0001). Data is presented as mean ± SD.

### Treatment of miR-9-5p antagomiRs altered time from peak to baseline in control iPSC-CM

To determine whether miR-9-5p downregulation in healthy control iPSC-CMs promotes contractile changes consistent with hypercontractility observed in HCM, we assessed contractile behaviour in miR-9-5p antagomiR-treated control iPSC-CM using PIV. In control iPSC-CMs (Figure 9A), treatment with miR-9-5p antagomiRs did not significantly affect maximum displacement during peak contraction (Figure 9B), contraction cycle length (Figure 9C), or time to peak contraction (Figure 9D), compared to the No Treatment group. Nevertheless, treated cells exhibited an increasing trend in these parameters with higher antagomiR doses, resembling the contractile phenotype observed in untreated *MYBPC3* mutant iPSC-CMs. A significant decrease in relaxation time was observed in control iPSC-CMs treated with 100 nM miR-9-5p antagomiRs (740 ± 340 ms) compared to No Treatment (1400 ± 480 ms; p = 0.0449). Additionally, cells treated with 100 nM showed a significantly shorter relaxation compared to those treated with 50 nM (800 ± 320 ms; p = 0.0182) (Figure 9E). Together, these findings suggest that miR-9-5p downregulation in control iPSC-CMs induces emerging, dose-dependent shifts in contractile kinetics toward a hypercontractile, disease-like phenotype characteristic of HCM, even in the absence of statistically significant changes in peak contraction magnitude.

**Figure 9.**
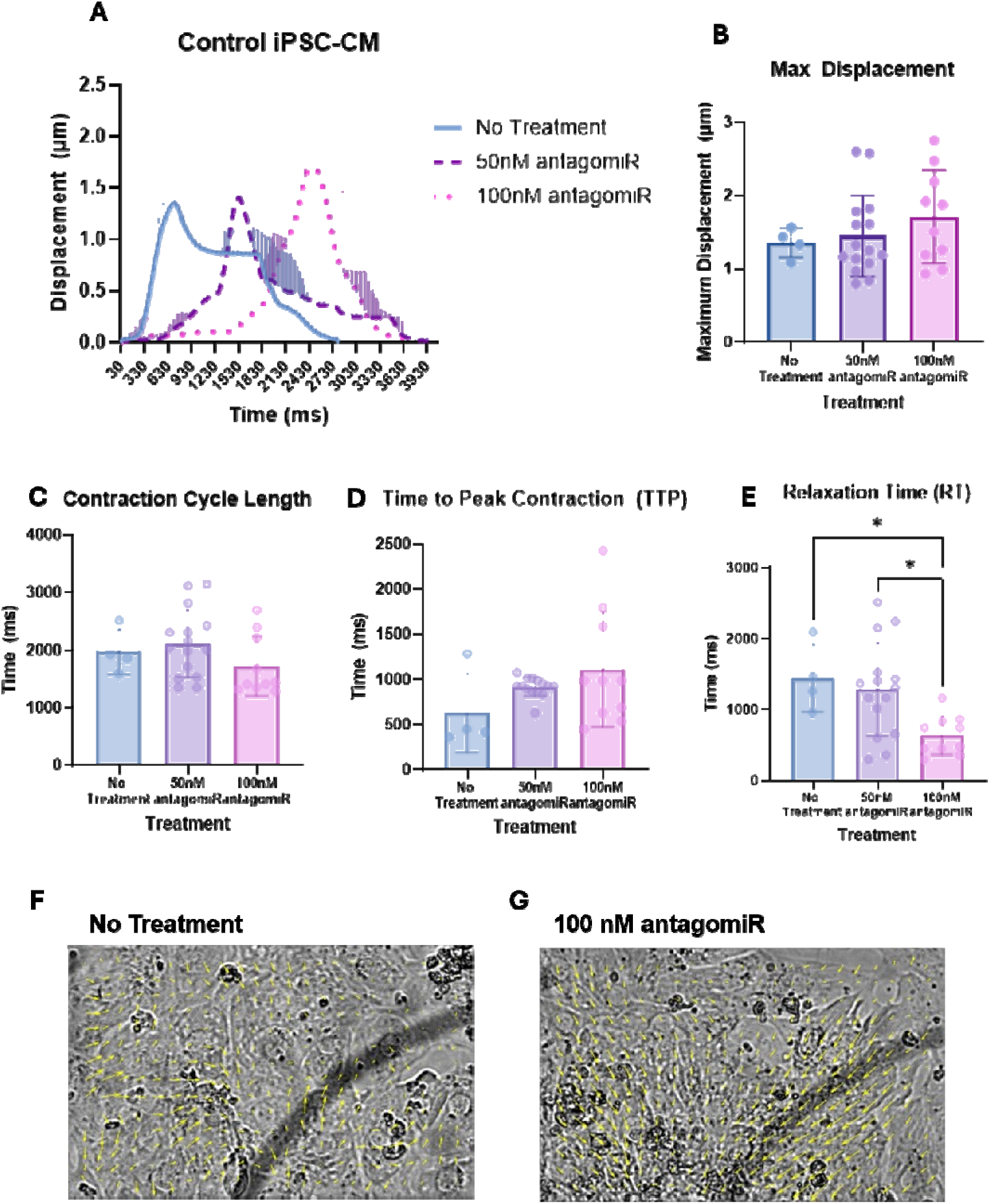
Contractility of control iPSC-CM observed using Particle Image Velocimetry (PIV). Images were obtained using phase contrast microscopy, and particle displacement magnitude (µm) was measured over time (ms). **(A)** Representative contraction curves of control iPSC-CMs: No Treatment (n = 4) or treated with 50 nM (n = 15) or 100 nM (n = 10) of miR-9-5p antagomiRs. Quantification of contractility parameters in control iPSC-CM: **(B)** Maximum displacement, **(C)** Contraction cycle length, **(D)** Time to peak contraction (TTP) **(E)** Relaxation time. **(F)** Vector plot arrows (in yellow) depicting contraction direction of untreated control iPSC-CM and **(G)** control iPSC-CM treated with 100 nM of miR-9-5p antagomiRs. Videos can be accessed at https://doi.org/10.5683/SP3/GUBIRH. Statistical analysis was performed using One-way ANOVA with Tukey’s multiple comparisons test (* p ≤ 0.05). Data is presented as mean ± SD.

### Modulation of EGFR-TKI resistance pathway targets by alteration in miR-9-5p

To assess the effects of miR-9-5p upregulation on the EGFR-TKI resistance pathway, total RNA was isolated from *MYBPC3* mutant iPSC-CMs treated with miR-9-5p mimics, followed by RT-qPCR analysis. This approach was used to gain mechanistic insight into signaling pathways that may be altered by miR-9-5p modulation in the context of HCM. Transcriptional analysis of genes within the EGFR-TKI resistance pathway revealed a coordinated shift toward reduced pro-growth signaling following miR-9-5p upregulation. Specifically, growth-promoting genes were significantly downregulated, including *PDGFRB* (fold change = 0.50, p = 0.0049 at 100 nM) and *IL6* (fold change = 0.55, p = 0.026 at 100 nM; 0.57, p = 0.0327 at 150 nM) (Figure 10A). *PDGFRB* is a known upstream activator of the PI3K/AKT and MAP/ERK pathways (Farooqi & Siddik, 2015), while *IL6* promotes cell growth through activation of the JAK/STAT pathway (Figure 10C) (Huang et al., 2022).

**Figure 10.**
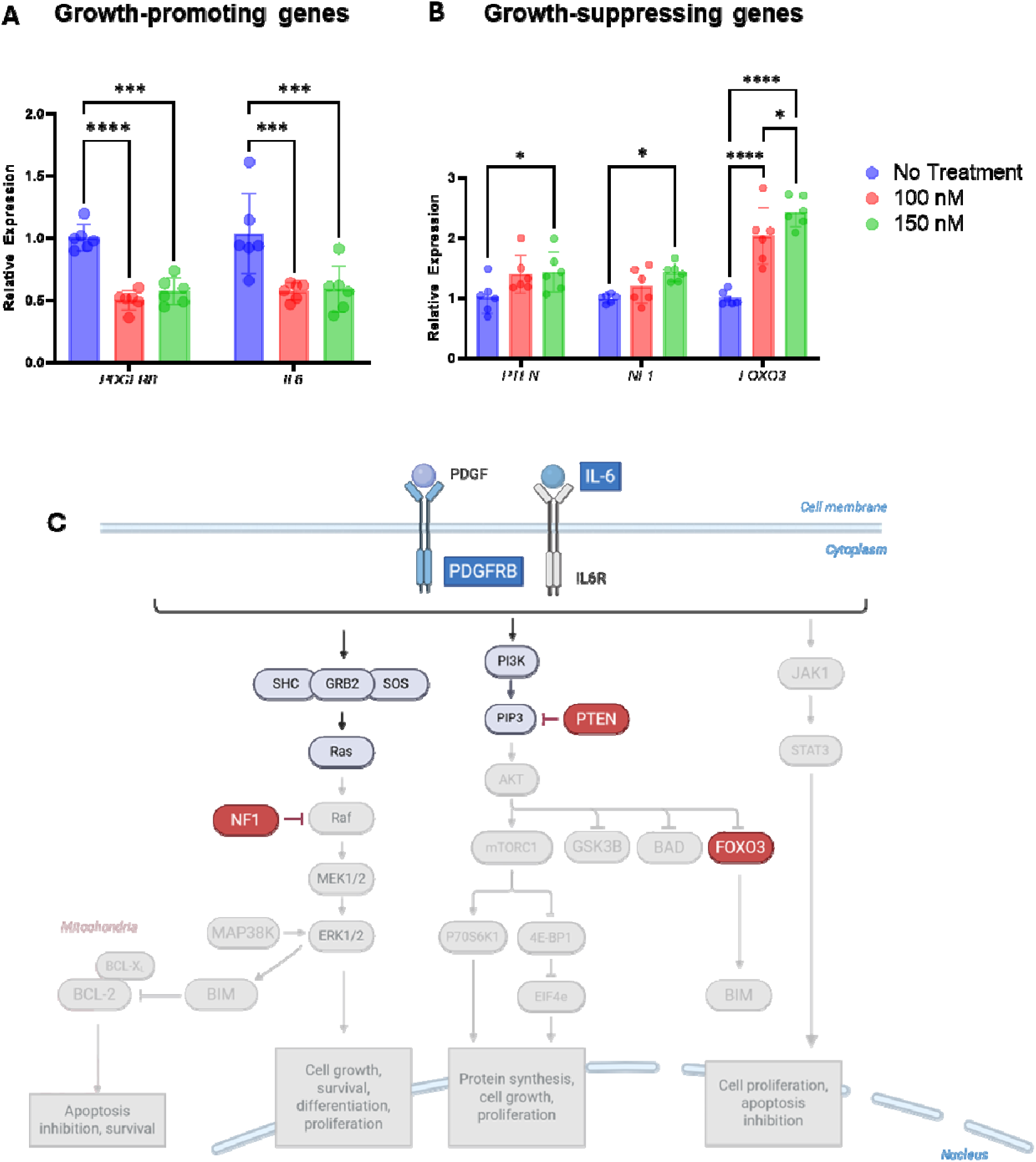
Modulation of EGFR-TKI resistance pathway in *MYBPC3* iPSC-CM treated with miR-9-5p mimics. **(A)** Transcriptional analysis of genes in EGFR-TKI pathway, including downregulation of growth-promoting genes *PDGFRB* and *IL6* and **(B)** upregulation of growth-suppressing genes *PTEN, NF1* and *FOXO3*. Relative gene expression was calculated using the 2^-ΔΔCt^ method and normalized to the 5S rRNA housekeeping gene set. **(C)** Proposed schematic of mechanism of action. Upregulation of miR-9-5p expression leads to inactivation of downstream pathways promoting cell growth and proliferation, including MAP/ERK, PI3K/AKT, and JAK/STAT.

Concurrently, several growth-suppressive genes were significantly upregulated, including *NF1* (fold change = 1.43, p = 0.0275 at 150 nM), *PTEN* (fold change = 1.40, p = 0.0368 at 150 nM), and *FOXO3* (fold change = 2.03, p < 0.0001 at 100 nM; 2.42, p < 0.0001 at 150 nM) (Figure 10B). *NF1* and *PTEN* negatively regulate MAP/ERK and PI3K/AKT signaling, respectively, while *FOXO3* represses transcription of genes involved in cellular growth and proliferation (Figure 10C).

Collectively, these transcriptional changes suggest that miR-9-5p upregulation attenuates downstream signaling through key nodes of the EGFR-TKI resistance pathway, including MAP/ERK, PI3K/AKT, and JAK/STAT. This coordinated suppression of pro-growth signaling provides a mechanistic basis for the observed reductions in cardiomyocyte size and altered cellular phenotypes in *MYBPC3* mutant iPSC-CMs treated with miR-9-5p mimics.

## Discussion

In this study, we investigated miRNA differential expression in HCM patients using an iPSC-CM model, with validation from myectomy tissues carrying mutations in the same genes. We identified significant downregulation of miR-9-5p in all mutant samples compared to healthy controls with validation through RT-qPCR in *MYBPC3* iPSC-CM, *MYH7* iPSC-CM, and myectomy tissues. Pathway analysis of genes regulated by miR-9-5p revealed significant enrichment of the EGFR tyrosine kinase inhibitor resistance pathway. Treatment with miR-9-5p mimics in *MYBPC3* mutant iPSC-CMs led to a significant reduction in CM size, changes in cell morphology, and increased nuclei and cell density. Upregulating miR-9-5p expression using mimics also modulated contractility, as indicated by decreased maximum displacement magnitude, shorter contraction duration, and reduced time from baseline to peak contraction. Conversely, inhibition of miR-9-5p expression in healthy control iPSC-CM using antagomiRs led to significantly increased CM surface area compared to the No Treatment and Control miRNA-treated groups. AntagomiR-treatment also led to decreases in CM circularity and roundness and led to a decrease in nuclei and cell density. Treatment of miR-9-5p antagomiRs had minimal effect on contractile parameters in healthy control iPSC-CM, except for reduced time from peak to baseline in antagomiR-treated groups. Together, these findings suggest that miR-9-5p plays a role in regulating both cardiomyocyte hypertrophy, proliferation, and contraction dynamics in iPSC-CM, and could be a potential target in mitigating disease phenotypes in HCM.

Previous studies have primarily discussed miR-9-5p in the context of cancer (G. Li et al., 2017; M. Wang et al., 2019; Zhu et al., 2021) and schizophrenia (Camkurt et al., 2016; Fu et al., 2023; Topol et al., 2016). In patients with carotid artery stenosis (CAS), miR-9-5p was significantly downregulated in asymptomatic cases and identified as a factor closely associated with cerebrovascular events, including strokes (Liu et al., 2021). In diabetic cardiomyopathy, miR-9-5p downregulation in diabetic mouse hearts was linked to cardiac fibrosis regulation via the circRNA_012164-miR-9-5p axis (Wang et al., 2024). Additionally, miR-9-5p plays a role in ischemic heart injury through the lncRNA-TUG1-miR-9-5p-KLF5 pathway, which regulates cardiomyocyte apoptosis (Yang et al., 2019). While the specific role of miR-9-5p in HCM has never been investigated, there have been associations of cardiac hypertrophy and miR-9, a closely related miRNA. miR-9 has been implicated in cardiac hypertrophy by modulating NFATc3 and myocardin in rat CM. Stimulating hypertrophy using isoproterenol and aldosterone significantly decreased miR-9 levels, while administering miR-9 ameliorated cardiac hypertrophy and improved cardiac function (K. Wang et al., 2010). Our findings position miR-9-5p as a previously unrecognized regulator of HCM, implicating its downregulation as a contributor to pathological CM growth, hypercontractility, and dysregulated pro-growth signaling pathways that underlie disease remodeling.

Consistent with prior studies linking miR-9-5p to cardiovascular disease, we observed significant downregulation of miR-9-5p in iPSC-CM and myectomy tissues from HCM patients. Treatment with miR-9-5p mimics in *MYBPC3* mutant iPSC-CM, led to a reduction in cardiomyocyte area, suggesting that miR-9-5p plays a role in regulating cell size. Conversely, treatment of miR-9-5p inhibitors in the healthy control iPSC-CM led to a significant increase in cardiomyocyte size. Interestingly, our findings also showed that miR-9-5p upregulation may increase proliferation, as shown by increased nuclei and cell density in our miR-9-5p mimic-treated *MYBPC3* mutant iPSC-CM. Conversely, inhibiting miR-9-5p in healthy control iPSC-CM using antagomiRs decreased nuclei and cell density compared to non-treated or Control miRNA-treated iPSC-CM. In cancer studies, miR-9-5p has demonstrated context-dependent effects on proliferation: it inhibited cell growth via NF-κB activation in ovarian cancer (Guo et al., 2009), yet promoted growth and metastasis in non-small cell lung cancer by repressing *TGFBR2* (Li et al., 2017). However, across multiple cancer models, miR-9-5p is more consistently identified as a tumor suppressor that inhibits proliferation (Bandini et al., 2020; Guo et al., 2018; Zhang et al., 2019).

In the literature, iPSC-CMs are consistently described as having a fetal or immature phenotype characterized by a rounded morphology, whereas mature adult CM are elongated and rod-shaped (Wu et al., 2021). In this study, circularity and roundness were used to characterize changes in cell morphology; however, these measures should not be interpreted as direct indicators of CM maturation. Circularity quantified cell shape complexity, with a value of 1.0 representing a perfect circle and lower values indicating increasingly irregular or elongated boundaries, while roundness provided a measure of cell elongation. *MYBPC3* mutant iPSC-CMs treated with 250 nM miR-9-5p mimics exhibited a significant reduction in circularity compared with Control miRNA-treated iPSC-CMs, whereas roundness remained unchanged. Similarly, healthy control iPSC-CMs treated with 50 and 100 nM miR-9-5p antagomiRs exhibited significantly reduced circularity, while roundness was significantly decreased at 100 nM. Although decreased roundness and circularity could be consistent with changes in cell morphology, the effect at this relatively high mimic concentration may alternatively reflect cellular stress or cytotoxicity rather than a maturation-associated phenotype. In particular, excessive delivery of exogenous miRNA may disrupt the endogenous miRNA regulatory machinery through increased loading of miRNAs into Argonaute-containing complexes, potentially resulting in off-target effects and cellular toxicity (Grimm et al., 2010). This possibility is particularly relevant at higher treatment concentrations and should be considered when interpreting the morphological changes observed following miR-9-5p overexpression.

In addition to altering CM morphology, treatment with miR-9-5p mimics in *MYBPC3* iPSC-CMs affected contractile behavior, reducing contraction magnitude, contraction duration, and the time from baseline to peak. Our model recapitulated hypercontractility through increased contraction magnitude, consistent with early cardiac features observed in HCM (Toepfer et al., 2019). Treatment with miR-9-5p mimics significantly attenuated contraction magnitude, total contraction duration, and time to peak, suggesting a potential modulatory effect on the hypercontractile phenotype. While there is limited literature on miR-9, a related miRNA, in the context of cardiac contractility, studies have demonstrated a similar effect in smooth muscle cells. In a study by Luo *et al*., administering miR-9 mimics decreased contractility, while miR-9 inhibitors showed higher contractility, suggesting that miR-9 weakened contractility in smooth muscle cells (Luo et al., 2016). A recent study identified dysregulated kinase inhibitors in *MYBPC3* mouse models that may contribute to impaired relaxation in HCM (Algül et al., 2023). Phosphoproteomic profiling of HCM patient myocardial tissues also revealed increased activation of six proteins within the EGFR/IGFR1 cluster, suggesting that EGFR dysregulation may contribute to contractile dysfunction in HCM (Algül et al., 2023). Additionally, in tissue-engineered iPSC-CM with *MYH7* mutations, EGFR was implicated in pathological phenotypes of HCM, and blocking EGFR signaling mitigated fibrotic and contractile abnormalities (Ewoldt et al., 2024).

In *MYBPC3* iPSC-CM treated with miR-9-5p mimics, there was a significant decrease in growth-promoting genes *PDGFRB* and *IL6,* as well as an increase in expression of growth-suppressing genes *NF1, PTEN,* and *FOXO3*. Overall, these results identify miR-9-5p as a central regulator of cell proliferation and growth in iPSC-CMs, with its transcriptional effects shaped by both dosage and cellular context. While transcriptional data and modulation of the EGFR-TKI resistance pathway provides an initial understanding of how miR-9-5p alters gene networks in cardiomyocytes, miRNAs exert much of their regulatory influence post-transcriptionally. mRNA abundance alone may therefore underestimate the full extent of miR-9-5p-mediated effects. To better define the downstream signaling consequences, we will perform phospho-proteomic analyses aimed at identifying the specific pathways and kinase activation events triggered by miR-9-5p modulation. These studies will clarify how transcriptional changes translate into functional signaling outputs and will help map the mechanisms through which miR-9-5p influences cardiomyocyte stress responses.

## Limitations

While this study establishes a clear association between miR-9-5p and cardiomyocyte remodeling, several limitations should be noted. The 2D iPSC-CM monolayer system offers advantages as a controlled and reproducible platform for investigating cellular responses to miRNA modulation. Its flat growth surface enables uniform exposure to treatments, high-resolution imaging, and precise quantification of morphological parameters, while remaining technically straightforward and cost-effective. However, this system does not fully recapitulate the three-dimensional architecture, mechanical cues, or cellular heterogeneity of native myocardial tissue. Additionally, iPSC-CMs in 2D culture remain structurally and functionally immature, exhibiting a fetal-like phenotype that may not accurately reflect adult cardiomyocyte behavior (Karakikes et al., 2015). However, despite differences between iPSC-CM and myectomy tissues, we consistently identified miR-9-5p as significantly downregulated, suggesting its critical role in development of HCM. Additionally, during the development of our in vitro model, *MYH7* iPSC colonies showed a tendency for spontaneous differentiation, and successfully differentiated iPSC-CMs often detached from the culture substrate. Due to these challenges, we conducted transfection experiments using *MYBPC3* mutant and control iPSC-CMs, which demonstrated greater stability. Future studies could involve conducting miR-9-5p transfection experiments in *MYH7* iPSC-CM lines to validate findings. Contractility measurements were derived from spontaneously beating cells rather than those paced by electrophysiological stimulation, limiting their physiological relevance. Variability among iPSC lines and differences in transfection efficiency may also contribute to experimental heterogeneity. Finally, the study focused on short-term responses to miR-9-5p modulation; longer-term effects on cellular maturation and function remain to be explored. Future studies should validate these findings in 3D engineered cardiac tissues, incorporate electrical pacing to better mimic in vivo physiology, and investigate downstream gene targets to delineate the precise mechanisms of miR-9-5p regulation.

## Conclusion

This study is the first to investigate miRNA expression in iPSC-CM with validation in myectomy tissues from HCM patients carrying *MYBPC3* and *MYH7* mutations. Using microarray analysis and RT-qPCR validation, we identified miR-9-5p as significantly downregulated in HCM compared to healthy controls. Pathway analysis revealed that genes regulated by miR-9-5p are involved in the EGFR tyrosine kinase inhibitor resistance pathway, previously implicated in dysregulated contractility and cardiomyocyte apoptosis in HCM. Further, treatment using miR-9-5p mimics led to decrease in cardiomyocyte size and contraction. Understanding how miR-9-5p contributes to cellular remodeling may provide new insights into the molecular basis of HCM and identify possible therapeutic avenues for mitigating pathological hypertrophy and hypercontractility.

## Materials and Methods

### Total RNA isolation from iPSC-CM and myectomy tissue

iPSCs were derived from blood of patients with HCM harboring the *MYBPC3* and *MYH7* mutation, and their respective family healthy family members and differentiated into cardiomyocytes by the Centre for Commercialization of Regenerative Medicine (CCRM) using a protocol by Lian and colleagues (Lian et al., 2013), and an enrichment method based on Sharun and colleagues (Sharma et al., 2015). iPSCs were differentiated into CM and frozen in stocks on post-differentiation Day 19. Total RNA was extracted from frozen iPSC-CM vials (∼2.0E06 cells) using the TRIzol method. Myectomy tissue samples were collected from patients with HCM harboring mutations in *MYBPC3* and *MYH7* and presented clinical symptoms, as well as healthy donors whose hearts were not eligible for transplant. Total RNA was isolated from <100 mg of myocardial tissue using the TRIzol method. RNA quality was measured from all samples using a Bioanalyzer (Agilent, 2100 Bioanalyzer).

### NanoString microarray analysis

A total of 12 uL of extracted RNA was used for NanoString microarray analysis, which involves a panel of 799 human miRNAs. Raw counts were processed using blank lanes to remove background noise and normalized using CodeSet Content Normalization of Top 100 counts. All raw data processing was conducted using nSolver (v.4.0) and miRNA counts were converted to fold changes between mutant and control lines.

### miRNA validation of microarray using reverse transcription quantitative polymerase chain reaction

The miRCURY LNA RT Kit (Qiagen, 339340) was used to perform reverse transcription of miRNA, we used according to manufacturer’s instructions using 500 ng of total RNA and conducted reverse transcription using Eppendorf Mastercycler (EP6301000012). cDNA was diluted 1:20 in nuclease-free water, and quantitative polymerase chain reaction (qPCR) was conducted using miRCURY LNA miRNA PCR Assays (Qiagen, 339306) according to manufacturer’s instructions. Normalized expression of miRNAs was determined using the 2^-ΔΔCt^ method and normalized using the 5S rRNA control primer set (V00589). All statistical analysis between mutant and control groups were conducted using Student’s t-test (Graphpad Prism, v.10.4.1).

### GO Term and KEGG pathway analysis using g:Profiler

Using miRTarBase (v.9.0), we generated a list of genes targeted by hsa-miR-9-5p (referred to as miR-9-5p) that were experimentally validated using strong methods: luciferase reporter assay, western blot, and quantitative polymerase chain reaction (Supplementary Table 2). An interactome was created via STRING (v.12.0) to visualize gene-gene interaction. Functional enrichment analysis was conducted using g:Profiler. P values of significantly enriched Gene Ontology (GO) and Kyoto Encyclopedia of Genes and Genomes (KEGG) terms were adjusted using the g:SCS algorithm, α = 0.05.

### iPSC Maintenance and iPSC-CM Differentiation for Transfection Experiments

Induced pluripotent stem cells (iPSCs) were seeded in Geltrex LDEV-Free Reduced Growth Factor Basement Membrane Matrix (ThermoFisher Scientific, A1413201) and maintained in mTeSR1 (STEMCELL Technologies, 85850). iPSCs were dissociated into single cells using TrypLE Select Enzyme 1X (ThermoFisher Scientific, 12563011) and re-plated in Matrigel (Corning, 354277)-coated plates at a density of 2.0×10^5^ cells/cm^2^ and differentiated into cardiomyocytes using STEMdiff Ventricular Cardiomyocyte Differentiation Kit (STEMCELL Technologies, 05010) and maintained in Cardiomyocyte Maintenance Media (STEMCELL Technologies, 05020).

### Transfection Experiments

miRIDIAN microRNA human hsa-miR-9-5p mimic (Horizon Discovery, C-300618-05-0005), miRIDIAN microRNA Human hsa-miR-9-5p antagomiR (Horizon Discovery, IH-300618-07-0005), and miRIDIAN microRNA hairpin inhibitor negative control #1 (Horizon Discovery, IN-001005-01-20) were resuspended in 1x siRNA Buffer (diluted from 5x siRNA Buffer (Horizon Discovery, B-002000-UB-100)) and stored in 10 μM stock solutions. On Day 19 of the cardiomyocyte differentiation protocol, iPSC-cardiomyocytes (CM) were digested in Collagenase, Type 2 (Cedarlane, LS004174) in Hank’s buffered saline solution for 1hr and dissociated into single cells using TrypLE Select Enzyme 1X (Thermofisher Scientific) for 20 mins. Single cell iPSC-CM were re-plated in Matrigel-coated plates at a seeding density of 2.0×10^5^ cells/cm^2^ for transfection experiments. On Day 20, *MYBPC3* mutant iPSC-CM were treated with 25 nM, 50 nM, 100 nM and 150 nM of miR-9-5p mimics, and healthy control iPSC-CM were treated with 25 nM, 50 nM, 75 nM and 100 nM of miR-9-5p antagomiRs using *Trans*IT-TKO Transfection Reagent (Mirus Bio, MIR 2154). Control miRNA conditions include 50 nM of the miRIDIAN microRNA hairpin inhibitor negative control #1. iPSC-CM were visualized at 24h post-transfection to ensure high efficiency (Supplementary Figure 2,3).Conditions for miRNA transfection with *Trans*IT-TKO Transfection Reagent are outlined below.

| Culture Vessel | 24-well plate |  | 12-well plate |  |
| --- | --- | --- | --- | --- |
| Surface Area | 1.9 cm<sup>2</sup> |  | 3.8 cm<sup>2</sup> |  |
| Cardiomyocyte<br>Maintenance<br>Medium | 0.5 mL |  | 1.0 mL |  |
| Serum-free<br>medium | 50 µL (varies depending on<br>desired final miRNA<br>concentration) |  | 100 µL (varies depending on<br>desired final miRNA<br>concentration) |  |
| *TransIT*-TKO<br>Reagent | 1 µL |  | 2.5 µL |  |
| miRNA (10µM<br>stock) | Final<br>Concentration | miRNA<br>amount | Final<br>Concentration | miRNA<br>amount |
|  | 25 nM | 1.4 µL | 25 nM | 2.8 µL |
|  | 50 nM | 2.8 µL | 50 nM | 4.2 µL |
|  | 75 nM | 4.2 µL | 75 nM | 5.6 µL |
|  | 100 nM | 5.6 µL | 100 nM | 8.4 µL |
|  | 150 nM | 8.4 µL | 150 nM | 11.2 µL |

### Immunofluorescence and morphology assessments

iPSC-CM were fixed with 2% paraformaldehyde for 10 mins at room temperature, then 90% ice-cold methanol for 10 mins. Cells were washed using PBS and permeabilized using permeabilization buffer (0.5% Triton X-100, 0.2% Tween-20 in PBS) for 30 mins at 4 °C. Blocking buffer (5% FBS in permeabilization buffer) was then added and incubated for 1 hr at room temperature. Cells were incubated with primary antibodies (α-actinin 1:500 (Abcam, ab9465), Connexin-43 1:500 (Abcam, ab217676)) in blocking buffer overnight at 4 °C on a slow rocker. Fluorophore-conjugated secondary antibody staining (anti-rabbit Alexa 488 or anti-mouse Alexa 594, 1:500 (Molecular Probes)) was conducted on a slow rocker at room temperature for 1 hr. Nuclear counterstaining was performed using 5 μg/mL DAPI at room temperature for 15 mins. Images were acquired using an Olympus FV3000 confocal microscope with 405, 488, and 561 nm lasers. Cardiomyocyte morphology was quantified using the *Measure* function in ImageJ (1.54p). Individual cells were outlined manually based on α-actinin and connexin-43 staining to capture cell boundaries. Areas with overlapping iPSC-CMs were excluded from measurements, and only cardiomyocytes with distinct borders were used for analysis. For each condition, multiple iPSC-CM were analyzed from four to nine images across two experimental replicates. Outlier analysis was conducted using the ROUT method (Q = 1 %) (Graphpad, Prism), and outliers identified were excluded before conducting statistical analysis between treatment groups using one-way ANOVA with Tukey’s multiple comparisons test (Graphpad, Prism). For each cell, Circularity and Roundness were extracted as shape descriptors generated by ImageJ. Circularity was calculated, where a value of 1.0 indicates a perfect circle and values approaching 0 indicate increasingly irregular shapes. Roundness, defined as the ratio of the minor to major axis length derived from the best-fit ellipse, reflects how close the cell shape is to a circle independent of area. A value of 1.0 indicates a perfect circle while values approaching 0 indicate more elongated or polarized polygons. Examples can be found in table below.

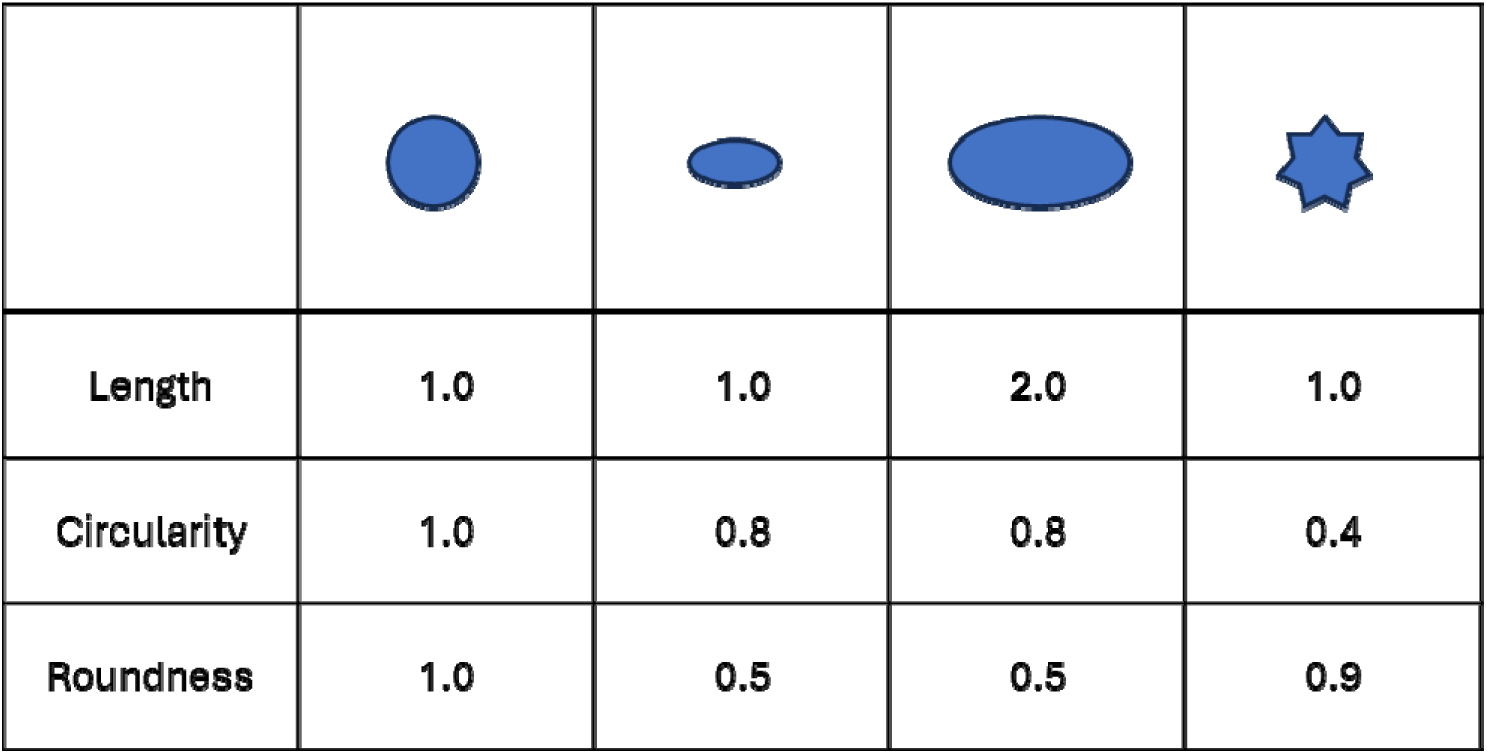

To calculate nuclear density, total number of nuclei was divided over total analyzed cardiomyocyte area, as reported by α-actinin expression. The number of multinucleated cells was derived from selected cardiomyocytes with distinct borders in each image.

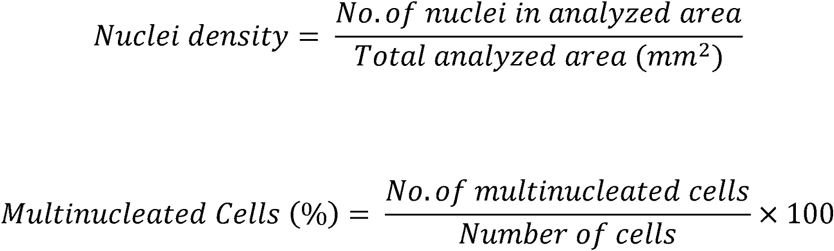

To estimate cell density while accounting for multinucleation, I used the following approximation to estimate the number of cells in each image assuming an average of two nuclei per multinucleated cell:

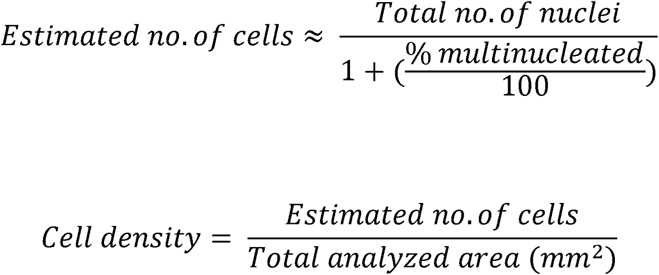

### Particle Displacement Velocimetry Image Acquisition and Analysis

Phase contrast images were acquired using an Olympus Epifluorescence Microscope (BX53) and BlackFly camera and micromanager Multi-D Acquisition (v.1.4) at 33.3 frames per second for 400 frames at 40x magnification. Images were uploaded onto MATLAB R2024a (24.1) PIVlab 2.63. Images were pre-processed with Enable Clache [50], and floating cells or debris were masked from PIV analysis. PIV settings were as follows: Pass 1 Interrogation Area: [128], Step: [64]. Pass 2 Interrogation Area: [80], Step: [40]. All results were transferred onto MATLAB workspace, and raw displacement magnitudes were exported. Contractility metrics, measured using the total average displacements of particles in the series of images, were calculated using the following formula:

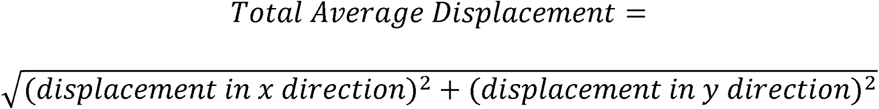

## Acknowledgements

Research in our laboratories on investigation of miRNA expression in hypertrophic cardiomyopathy has been funded by a Collaborative Health Research Project to Drs. Filio Billia and Craig Simmons from the Natural Sciences and Research Council of Canada (CHRPJ 5083) and Canadian Institutes of Health Research (CIHR; CPG-151946). Doris Adao is supported by a Queen Elizabeth II/Heart and Stroke Foundation of Ontario Graduate Scholarship in Science and Technology Award, and the Ted Rogers Centre for Heart Research Education Fund. Patient variables were collected by T. Hussain, S. Gramolini and J. Halpern.

## CRediT author statement

**Doris M.T. Adao:** Conceptualization, Methodology, Formal Analysis, Investigation, Writing – Original Draft, Visualization; **Heta Lad:** Methodology, Formal Analysis; **Mingyi Tang:** Methodology, Formal Analysis; **Sophia G. Lopez:** Methodology, Investigation; **Craig A. Simmons:** Conceptualization, Writing, Visualization, Writing – Review & Editing; Supervision; Project Administration; Funding Acquisition; **Filio Billia:** Conceptualization, Writing, Visualization, Writing – Review & Editing; Supervision; Project Administration; Funding Acquisition.

## Notes

### Competing Interest Statement

The authors have declared no competing interest.

