## Supplemental data for "Transcriptomic analysis of human induced pluripotent stem cell cardiomyocytes and myocardial tissue reveals novel role for miR-9-5p in hypertrophic cardiomyopathy"

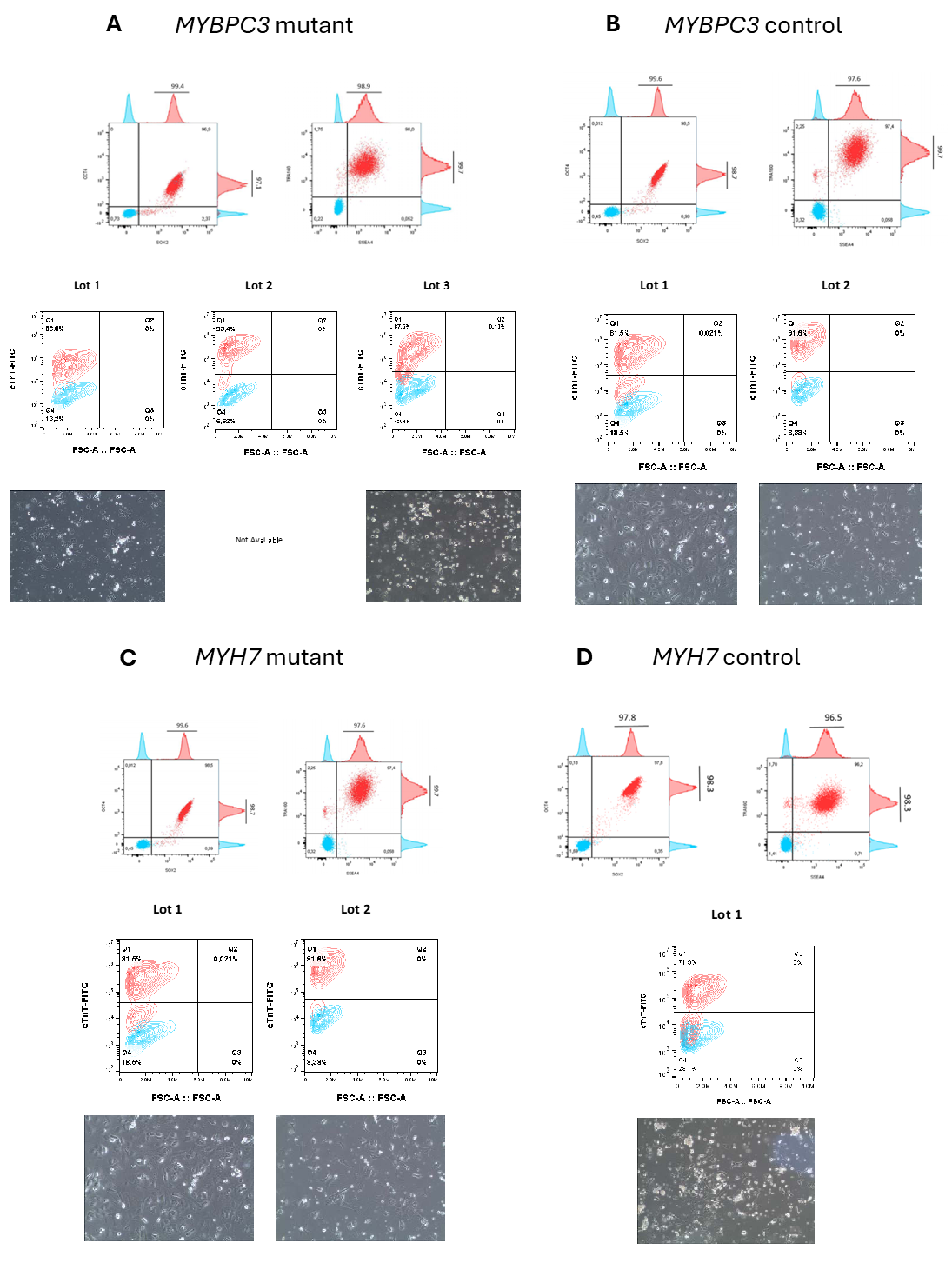

**Supplementary Figure 1.** Flow cytometry analysis and characterization conducted by CCRM for quality control of iPSCs (≥ 80% of population is positive for expression of surface markers (SSEA4, Tra-1-60), and intracellular marker (OCT4) and iPSC-cardiomyocytes (≥ 70% of population is positive for expression of intracellular marker (cardiac troponin (cTnT)) generated from various populations: **(A)** *MYBPC3* mutant, **(B)** *MYBPC3* control, **(C)** *MYH7* mutant, and **(D)** *MYH7* control.

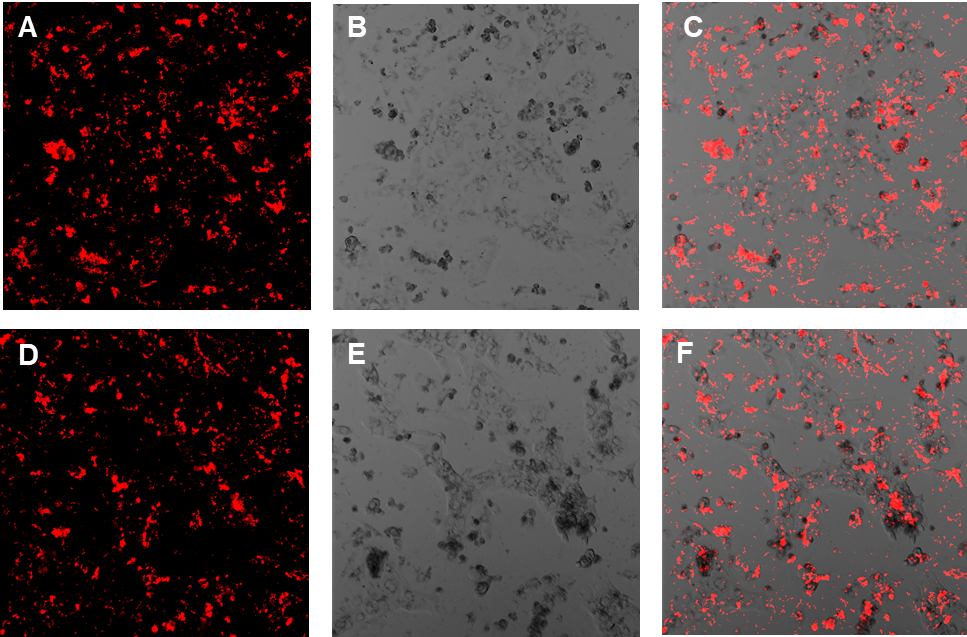

**Supplementary Figure 2.** Images of iPSC-CM 24-hours post-transfection to visualize transfection efficiency. Cells were plated onto 24-well plates and transfected with 50 nM of RFP using 1.0 μL of *TransIT*-TKO Transfection Reagent (Mirus Bio). **(A, D)** RFP only, captured using 561 nm lasers **(B, E)** iPSC-CM only, captured using Brightfield lens, **(C, F)** Merged. Images were captured using an Olympus FV3000 Confocal Microscope.

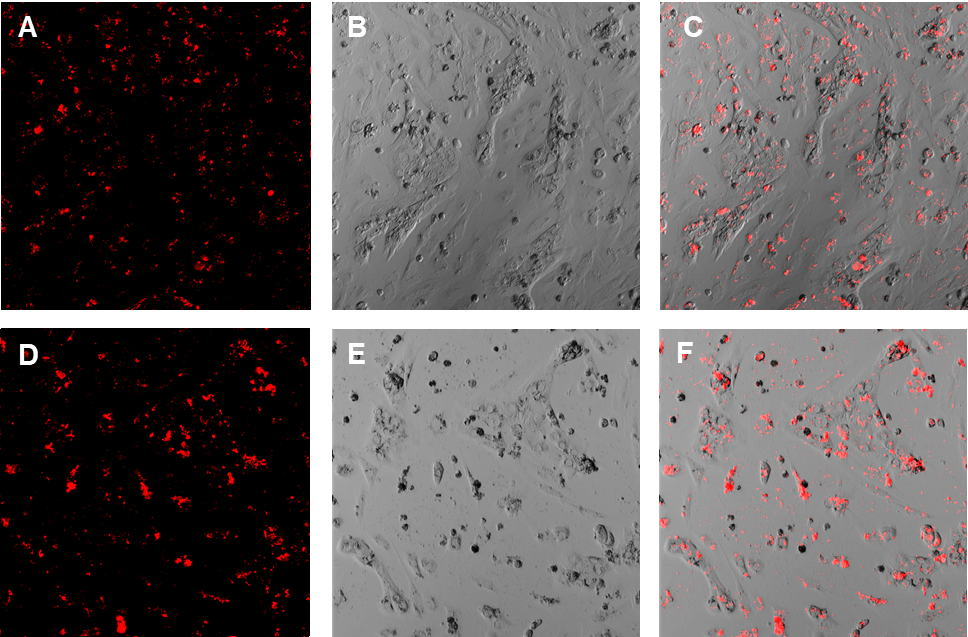

**Supplementary Figure 3.** Images of iPSC-CM 24-hours post-transfection to visualize transfection efficiency. Cells were plated onto 12-well plates and transfected with 50 nM of RFP using 2.5 μL of *TransIT*-TKO Transfection Reagent (Mirus Bio). **(A, D)** RFP only, captured using 561 nm lasers **(B, E)** iPSC-CM only, captured using Brightfield lens, **(C, F)** Merged. Images were captured using an Olympus FV3000 Confocal Microscope.

**Supplementary Table 2.** Fold-change expression and p-value of gene terms in EGFR-tyrosine kinase inhibitor resistance pathways, separated by sample type.

| Sample Type | Gene | Fold-change | p-value |
| --- | --- | --- | --- |
| *MYBPC3*  iPSC-CM | *FOXO3* | 1.52 | 0.026 |
|  | *NF1* | 1.53 | 0.0047 |
|  | *PTEN* | 1.91 | 0.015 |
|  | *GSK3B* | 2.05 | 0.022 |
|  | *PDGFRB* | 0.0054 | < 0.001 |
| *MYBPC3*  Myectomy tissue | *NF1* | 1.24 | 0.025 |
|  | *BCL2* | 3.06 | 0.018 |
|  | *FOXO3* | 0.64 | 0.022 |
|  | *GSK3B* | 0.82 | < 0.001 |
| *MYH7*  iPSC-CM | *PDGFRB* | 0.23 | < 0.001 |
|  | *NF1* | 0.75 | 0.026 |
|  | *BCL2* | 0.45 | < 0.001 |
|  | *BCL2L11* | 2.06 | < 0.001 |
| *MYH7*  Myectomy tissue | *FOXO3* | 0.61 | 0.022 |
|  | *JAK1* | 0.51 | 0.013 |
|  | *BCL2* | 3.57 | < 0.001 |
